# Vertical transmission of African-lineage Zika virus through the fetal membranes in a rhesus Macaque (*Macaca mulatta*) model

**DOI:** 10.1101/2023.03.13.532348

**Authors:** Michelle R. Koenig, Ann M. Mitzey, Xiankun Zeng, Leticia Reyes, Heather A. Simmons, Terry K. Morgan, Ellie K. Bohm, Julia C. Pritchard, Jenna A. Schmidt, Emily Ren, Fernanda Leyva Jaimes, Eva Winston, Puja Basu, Andrea M. Weiler, Thomas C. Friedrich, Matthew T. Aliota, Emma L. Mohr, Thaddeus G. Golos

## Abstract

Zika virus (ZIKV) can be transmitted vertically from mother to fetus during pregnancy, resulting in a range of outcomes, including severe birth defects and fetal/infant death. Potential pathways of vertical transmission *in utero* have been proposed but remain undefined. Identifying the timing and routes of vertical transmission of ZIKV may help us identify when interventions would be most effective. Furthermore, understanding what barriers ZIKV overcomes to effect vertical transmission may help improve models for evaluating infection by other pathogens during pregnancy. To determine the pathways of vertical transmission, we inoculated 12 pregnant rhesus macaques with an African-lineage ZIKV at gestational day 30 (term is 165 days). Eight pregnancies were surgically terminated at either seven or 14 days post-maternal infection. Maternal-fetal interface and fetal tissues and fluids were collected and evaluated with RT-qPCR, *in situ* hybridization for ZIKV RNA, immunohistochemistry, and plaque assays. Four additional pregnant macaques were inoculated and terminally perfused with 4% paraformaldehyde at three, six, nine, or ten days post-maternal inoculation. For these four cases, the entire fixed pregnant uterus was evaluated with *in situ* hybridization for ZIKV RNA. We determined that ZIKV can reach the MFI by six days post-infection and infect the fetus by ten days. Infection of the chorionic membrane and the extraembryonic coelomic fluid preceded infection of the fetus and the mesenchymal tissue of the placental villi. We did not find evidence to support a transplacental route of ZIKV vertical transmission via infection of syncytiotrophoblasts or villous cytotrophoblasts. The pattern of infection observed in the maternal-fetal interface provides evidence of vertical ZIKV transmission through the fetal membranes.

**Author’s Summary:** Zika virus (ZIKV) can be vertically transmitted from mother to fetus during pregnancy resulting in adverse pregnancy outcomes. For vertical transmission to occur, ZIKV must overcome the barriers of the maternal-fetal interface, yet the exact pathway ZIKV takes remains undefined. The maternal-fetal interface consists of the maternal decidua, the placenta, and the fetal membranes. ZIKV could reach the fetus through the placenta if it can infect the layer of cells that are directly exposed to maternal blood. ZIKV could also reach the fetus by infecting the decidua and then the adjacent fetal membranes. To determine the pathways of ZIKV vertical transmission, we infected pregnant macaques and evaluated ZIKV burden in the maternal-fetal interface and fetus shortly after maternal infection. The pattern of infection observed suggests that ZIKV vertically transmits through the fetal membranes, not the placenta. This finding is significant because it challenges the assumption that vertical transmission occurs exclusively across the placenta. By including the fetal membranes in our models of vertical transmission, we can more accurately determine which pathogens can be vertically transmitted. Ultimately, this study demonstrates that fetal membranes are an essential barrier to pathogens that warrant further investigation.

## Introduction

Zika virus (ZIKV) infection during pregnancy can cause birth defects and adverse pregnancy outcomes. Epidemiological evidence suggests that 5–10% of ZIKV infections during pregnancy result in birth defects or abnormal fetal development. Birth defects resulting from congenital ZIKV infection include microcephaly, joint contractures, ocular defects, and hearing abnormalities, collectively known as Congenital ZIKV syndrome (CZS) [1–8]. Congenital infection/exposure to ZIKV can also result in other adverse pregnancy outcomes, including fetal death, miscarriage, and intrauterine growth restriction [1,3,9]. Currently, we do not know the pathway ZIKV infection takes to transverse the maternal-fetal interface (MFI) to reach the fetus in an infected pregnant person; however, potential routes have been proposed [10].

One potential route of ZIKV vertical transmission is through the placenta, or “transplacental.” In this vertical transmission route, ZIKV in the maternal blood would directly infect the syncytiotrophoblasts (STBs) in the placenta, the cells in contact with the maternal blood, to facilitate oxygen and nutrient exchange. ZIKV then goes on to infect the inner layer of villous cytotrophoblasts (CTBs) to reach the fetal blood vessels inside the villi. Evidence for the transplacental route of ZIKV vertical transmission is currently sparse in human cases of congenital ZIKV infection and non-human primate (NHP) *in vivo* studies. To our knowledge, no published results show STB infection in cases of human congenital ZIKV infection, and ZIKV has rarely been found in STBs in NHP studies [11–15]. Reports of the vulnerability of STBs and CTBs to infection have primarily been obtained from *ex vivo* or *in vitro* studies using both Asian and African-lineage ZIKV. *Ex vivo* and *in vitro* studies provide somewhat conflicting results. Some studies show that STBs are resistant to infection due to potent type III interferon responses [10,16,17]; however, other studies find that derived and differentiated STBs are susceptible [18–20]. This inconsistency may reflect the maturity of the derived STBs, which may be more representative of primitive STBs present during early implantation rather than those present in the more mature placenta [20]. Overall, *ex vivo* and *in vitro* data suggest that mature STBs are generally resistant to ZIKV infection but may be susceptible to ZIKV infection very early in pregnancy [20, 21].

The second proposed vertical transmission route bypasses the placenta by going around the placenta (“paraplacental”) and through the fetal membranes. The fetal membranes are comprised of two distinct membranes: the chorionic membrane and the amniotic membrane (i.e., amnion). In early pregnancy, these membranes are spatially separated. The chorionic membrane is the outer membrane that originates from the trophectoderm of the blastocyst. The amnion is the inner membrane and originates from the inner cell of the blastocyst and encapsulates the embryo/fetus. As the fetus grows, the amnion is pushed against the chorionic membrane, and the membranes fuse; thus, these membranes are commonly collectively referred to as the fetal membranes.

In a paraplacental route, ZIKV in the maternal blood would infect the decidua and spread to the adjacent chorionic membrane. Once the chorionic membrane is infected, the fetus is no longer protected by trophoblasts and ZIKV can, theoretically, easily spread to fetal vessels in the chorionic plate and into the amniotic fluid. Substantial evidence supports that the fetal membranes are susceptible to ZIKV infection [10,11,14,15,17,22–27]. The fetal membranes have been found to be infected in a human case of congenital ZIKV infection that resulted in a miscarriage [22]. The vulnerability of the fetal membranes has been demonstrated in a wide range of studies, including NHP studies and various *ex vivo* and *in vitro* studies [10,11,14,15,17,23–27]. Additionally, studies show that the fetal membranes are susceptible to both African and Asian-lineage ZIKV and remain susceptible throughout gestation [10,17,27].

Despite what we have learned regarding the susceptibility of key tissues to ZIKV, we have yet to determine if ZIKV is vertically transmitted through a transplacental route, paraplacental route, or a combination of both to reach the fetus. Determining the pathway of vertical transmission, the time of initial fetal infection and the development of fetal pathology are necessary to understand when therapeutic strategies may be administered to prevent adverse pregnancy outcomes. Furthermore, determining what cellular barriers ZIKV overcomes to reach the fetus will help us develop better models to evaluate the ability of other pathogens to transmit vertically during gestation. Past attempts to establish a macaque model to study the pathways of vertical transmission have been difficult due to high inter-animal variability or a low vertical transmission rate. Therefore, in this study, we used an African-lineage ZIKV strain (ZIKV-DAK) that is known to result in a high rate of vertical transmission [28]. To determine the pathway of ZIKV vertical transmission, we inoculated 12 pregnant rhesus macaques (*Macaca mulatta*) at gestational day (gd) 30 with ZIKV-DAK and performed timed pregnancy terminations shortly after maternal inoculation. The timing and pattern of virus appearance in the components of the MFI and the fetus suggest that ZIKV-DAK is vertically transmitted via the chorionic membrane, i.e., a paraplacental route.

## Results

### Subcutaneous inoculation with 10^4^ PFU of ZIKV-DAK resulted in productive infection and neutralizing antibody responses

Twelve pregnant rhesus macaques were subcutaneously inoculated with 10^4^ plaque forming units (PFU) of a Senegalese isolate of African-lineage ZIKV (ZIKV-DAK) at approximately gd 30 (Fig 1). To minimize the variability in vertical transmission, we inoculated all dams at approximately gd 30, during the late embryonic stage (term = gd 165). Inoculation at this gestational age has been shown to result in a high rate of vertical transmission [28]. Details on the gestational age of each dam at the time of inoculation can be found in S1 Table. Blood was collected from the dams prior to inoculation, during the first two weeks after inoculation, and intermittently up to 87 days post-infection (dpi) or until euthanasia. Physical examination of infected dams found none of the ZIKV-associated symptoms reported in human infections, such as rash or conjunctivitis following inoculation. This finding is consistent with previously reported studies of ZIKV-DAK infection in rhesus macaques [14,15,23].

**Fig 1.**
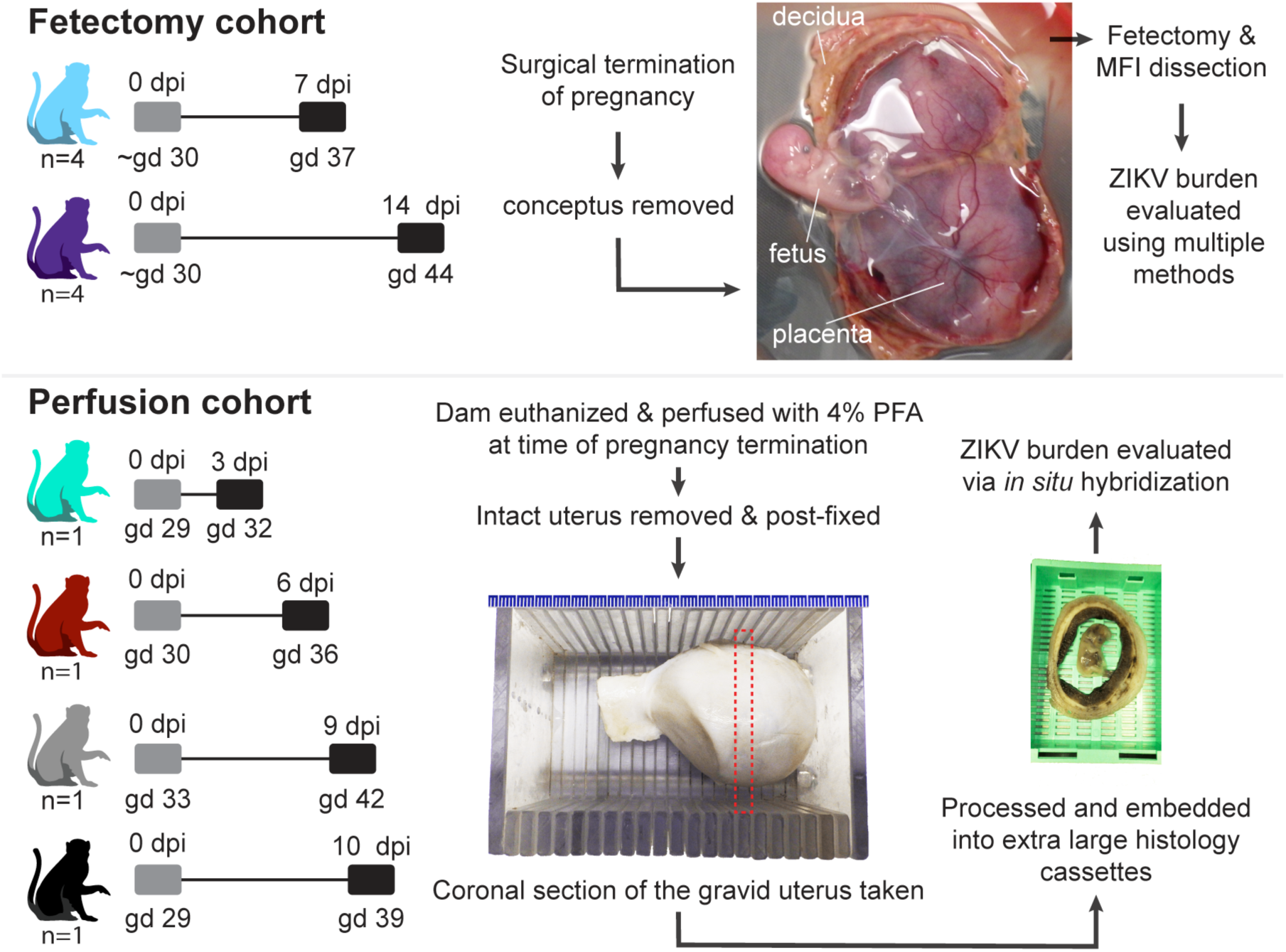
Experimental design. The pathway to vertical transmission was evaluated using two cohorts of pregnant rhesus macaques. The fetectomy cohort consisted of a total of eight dams that were inoculated with ZIKV-DAK at gestational day (gd) 30. These pregnancies were surgically terminated (four at 7 days post infection (dpi) and four at 14 dpi), the conceptus was removed and a fetectomy and maternal-fetal interface (MFI) dissection was performed to collect specimens that were evaluated for ZIKV burden using multiple methods. The perfusion cohort consisted of four dams that were inoculated with ZIKV-DAK at gd 30. At the time of pregnancy termination (either 3, 6, 9, or 10 dpi), the dams underwent terminal perfusion with 4% paraformaldehyde (PFA), and the gravid uterus was collected. The post-fixed uterus was sliced using a slicing box, creating consistent coronal sections of the entire uterus. Dashed line depicts a coronal section. These coronal sections were routinely processed and paraffin embedded using extra-large histology cassettes and viral burden in the coronal sections was evaluated using *in situ* hybridization. Gray boxes indicate the day of inoculation and black boxes indicate pregnancy termination.

This study consists of two cohorts of ZIKV-inoculated pregnant dams; eight dams in the “fetectomy cohort,” and four dams in the “perfusion cohort” (Fig 1). All twelve dams developed plasma viremia as expected (S1 Fig). To evaluate neutralizing antibody (nAb) responses to viral inoculation, we measured nAb titers in the serum prior to inoculation and either on the day of euthanasia or at approximately 28 dpi. Neither cohort of dams had detectable nAb prior to infection, confirming that these dams were immunologically naïve to ZIKV. All eight dams in the fetectomy cohort developed robust nAb following ZIKV inoculation as expected (S2 Fig). In the perfusion cohort, 10-1 had low levels of nAb at 10 dpi, while 09-1, 06-1, and 06-1 did not have detectable nAb levels at the time of euthanasia (S3 Fig). This result in the perfusion cohort is expected as these were euthanized three to nine days after infection before nAbs would likely develop.

### Evaluation of ZIKV-DAK burden in the MFI and fetus at 7 and 14 dpi

To determine the timing and pathway of ZIKV-DAK vertical transmission, we first evaluated the conceptus at seven and 14 dpi (Fig 1). In the fetectomy cohort, pregnancies were surgically terminated with collection of the conceptus, and sampling of the fetus and the MFI. Samples from the MFI and fetus were evaluated for ZIKV RNA using RT-qPCR and *in situ* hybridization (ISH). This study defines the MFI tissues as the decidua, the placenta, the chorionic membrane, and the amnion. Fluids from the conceptus were evaluated for ZIKV RNA using RT-qPCR and, when enough fluid was obtained, plaque assays were performed to test for infectious virus.

### ZIKV infects Extravillous trophoblasts (EVTs) at 7 dpi

Infection in the decidua was observed in all four pregnancies collected at 7 dpi (Fig 2). Evaluation of the decidua using ISH found ZIKV detected within a vessel occluded by extravillous trophoblasts (EVTs), forming an EVT’ plug’ that is normal at this gestational age [29] (Fig 3 A–C). Endovascular EVTs invade decidual spiral arteries to facilitate spiral artery remodeling, a process that is essential to a successful pregnancy [30–33]. The location of ZIKV as detected by ISH (Fig 3B) and the EVTs, as defined by cytokeratin expression (Fig 3C), suggests that the EVTs are infected with ZIKV. ZIKV was also detected in other decidual cells, including in macrophages (Fig 3D–F). ZIKV RNA was also seen at 7 dpi in the trophoblastic shell, the border zone of the placenta where EVTs anchor the macaque placenta to the uterus (Fig 3G, H). Immunohistochemistry (IHC) staining for cytokeratin suggested that the ZIKV RNA is detected within the EVTs of the trophoblastic shell (Fig 3G, H). These data indicate that both the endovascular EVTs and the EVTs in the trophoblastic shell are likely susceptible to ZIKV infection.

**Fig 2.**
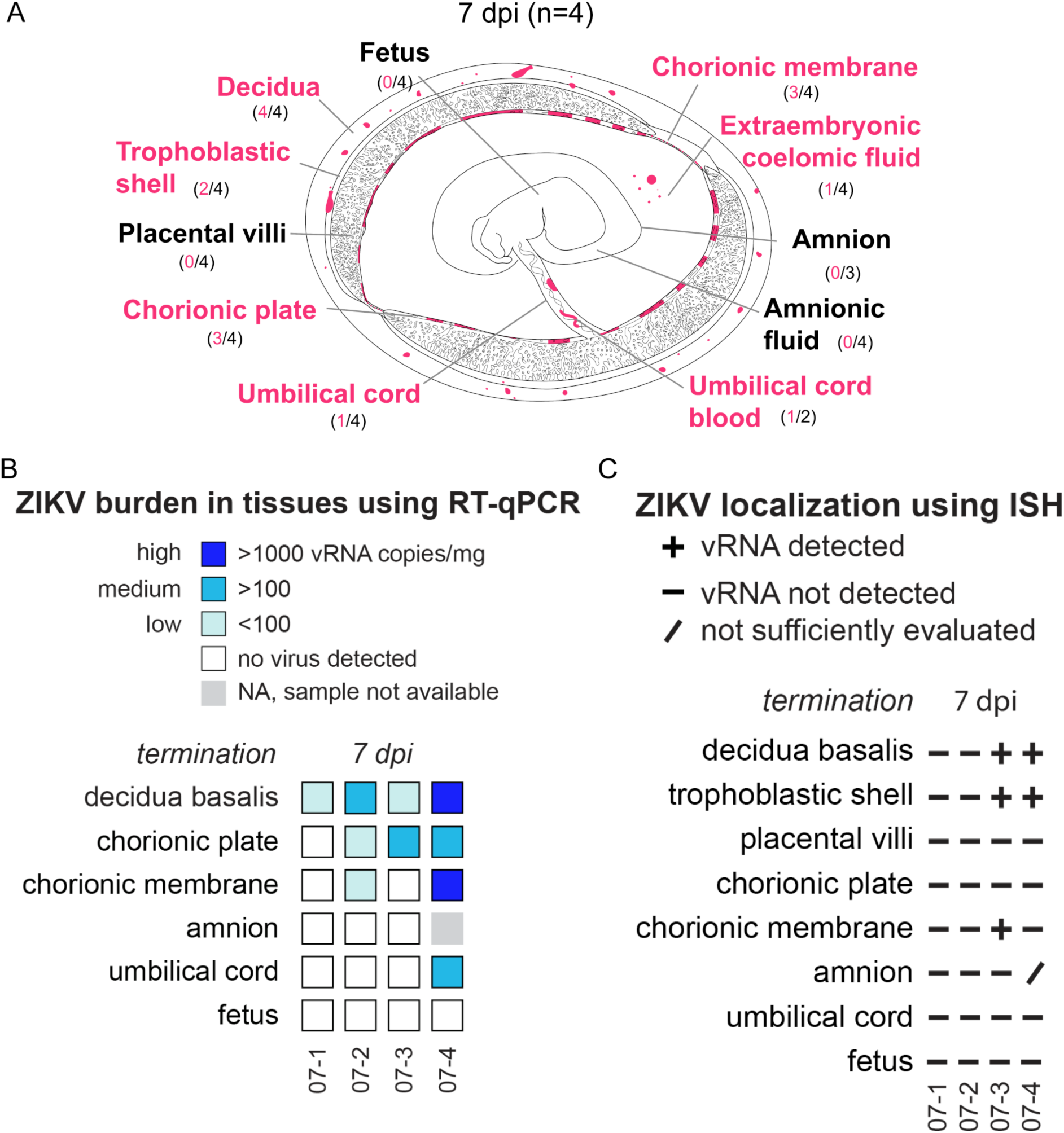
Summary of ZIKV viral burden at 7 dpi. (A) Artistic representation of ZIKV infection in the maternal-fetal interface at 7 days post infection (dpi). Pink tissue/structure names denote that ZIKV RNA was detected via ISH or RT-qPCR. The number of cases where the tissue had ZIKV RNA detected out of the number of cases in which tissue was evaluated is indicated below each tissue name. An artistic representation of the infection of each tissue is illustrated in pink in the figure. (B) Summary of the ZIKV RNA detected via RT-qPCR. The level of ZIKV burden is summarized as high (>1000 ZIKV RNA copies/mg of tissue), medium (>100 ZIKV RNA copies/mg), low (<100 copies/mg of tissue), no virus detected (below the limit of detection), or NA, sample not available. The theoretical limit of detection for all the tissues is 3 copies/mg. C) Summary of ZIKV RNA detected via ISH. ZIKV RNA was noted as present (+) or absent (-) in the indicated tissues/structures. In some instances, adequate histological samples were not obtained, and could not be properly evaluated (/). (B) (C) Individual animal IDs are listed on the bottom.

**Fig 3.**
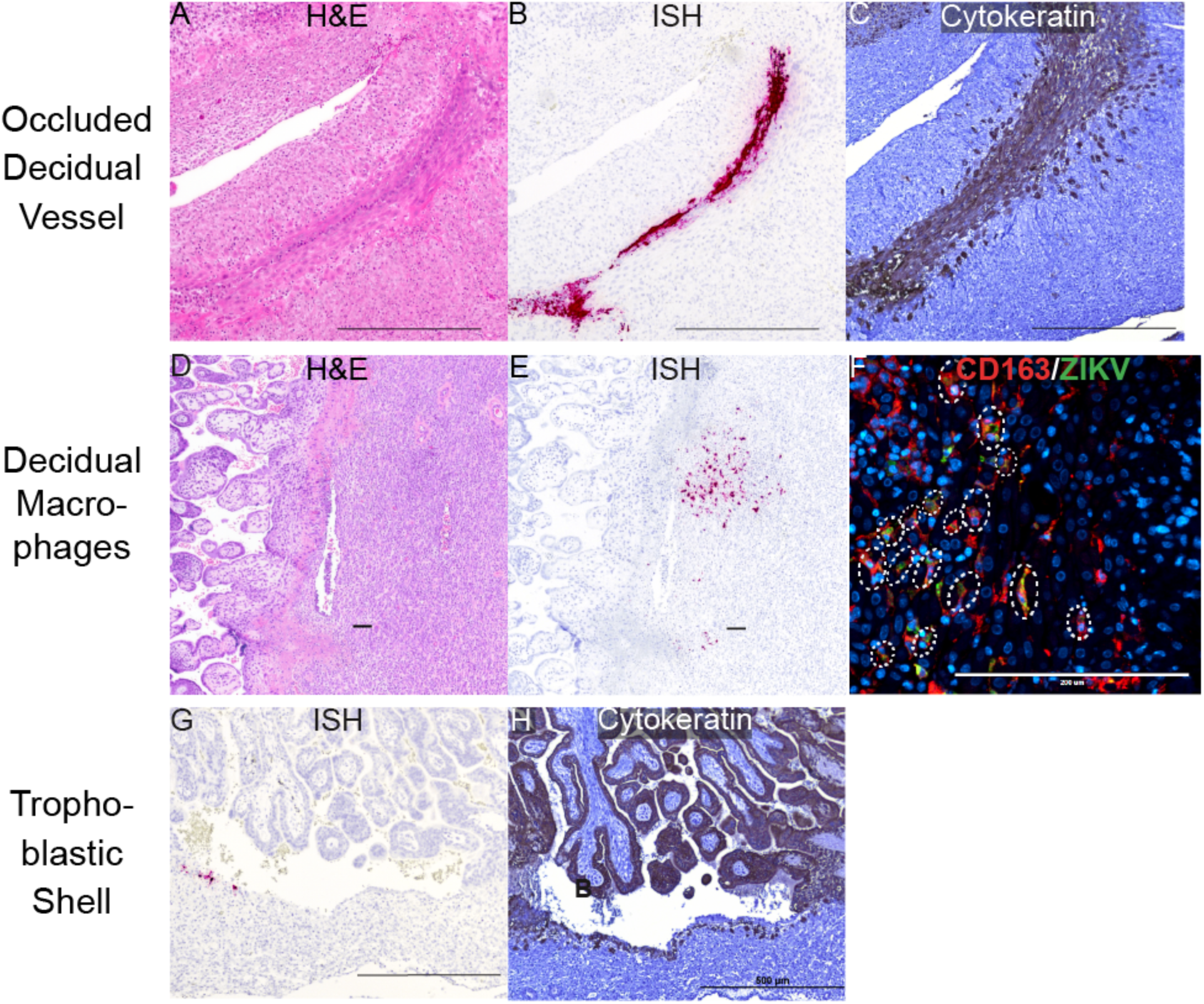
ZIKV infection in the decidua and trophoblastic shell at 7 dpi. ZIKV infection was seen in (A)–(C) a decidual vessel, (D)–(E) decidual macrophages, and (G)(H) in the trophoblastic shell. (A) H&E staining of the section shown in (B) Pink staining shows ZIKV RNA detected via ISH (C) IHC staining for cytokeratin. (A)–(C) Scale bar represents 500 µm. (D) H&E staining of the corresponding section shown in (E). (E) ISH (pink) staining showing ZIKV RNA in the decidua. (D)(E) Scale bars represent 100 µm. (F) IF staining for CD163 (red) to identify macrophages, ZIKV (green), and DAPI nuclear staining (blue). Colocalization of CD163, ZIKV, and DAPI is outlined by the dashed circles. The scale bar represents 200 µm. (G) Pink staining shows ZIKV RNA detected via ISH (H) IHC staining for cytokeratin, highlights the trophoblastic shell. (G)(H) Scale bars represent 500 µm.

### ZIKV infects the chorionic membrane and extraembryonic coelomic fluid but not the placental villi at 7 dpi

At 7 dpi, three out of four pregnancies had ZIKV RNA detected in the chorionic membrane by either RT-qPCR or ISH (Fig 2). The chorionic membrane consists of two layers: the trophoblastic and mesenchymal layers. ISH results show that ZIKV is in the trophoblast layer of the chorionic membrane (Fig 4). In early gestation, the space between the chorionic and amniotic membrane is filled with extraembryonic coelomic fluid (Fig 2A). Infectious ZIKV was detected in the extraembryonic coelomic fluid in one case, 7-3, at 7 dpi (Fig 5), ZIKV was not detected in the other three cases (S4A Fig). ZIKV RNA was also detected via RT-qPCR in the chorionic plate in three cases, and in one case in the umbilical cord at 7 dpi (Fig 2B). Because ZIKV RNA was only detected by RT-qPCR and not by ISH, we could not determine the cellular location of ZIKV in these tissues. ZIKV RNA was also seen in one case in the umbilical cord blood but there was not enough cord blood for a plaque assay (Fig 2, S4B Fig). No ZIKV RNA or infectious virus was detected above the limit of detection in the amnion, amniotic fluid, or placental villi at 7 dpi (Fig 2). Evidence of ZIKV RNA in the chorionic membrane, specifically in the trophoblasts of the chorionic membrane, and infectious virus in the extraembryonic coelomic fluid and the absence of ZIKV RNA in the placental villi demonstrates that ZIKV can bypass the placenta.

**Fig 4.**
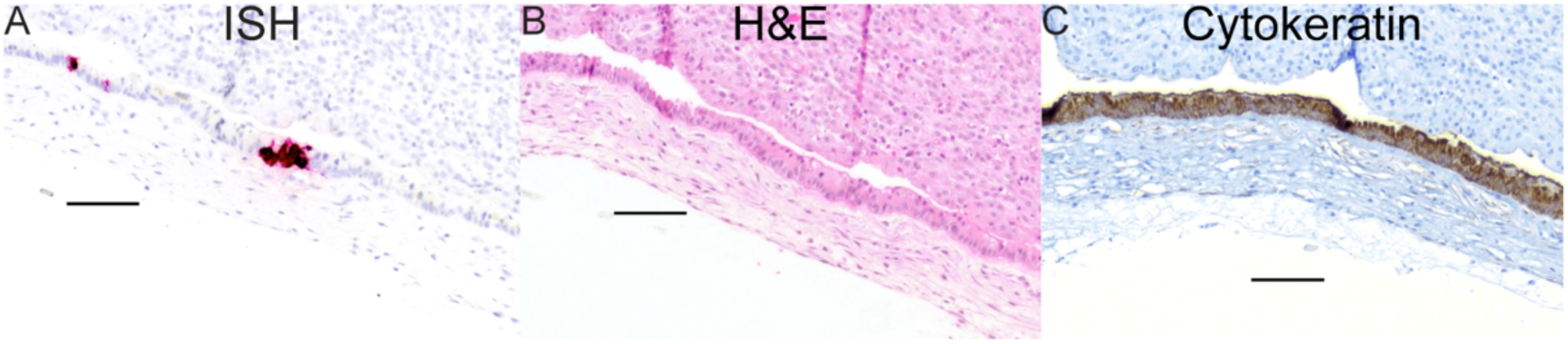
ZIKV infection of the chorionic membrane at 7 dpi. (A) Pink staining showing ZIKV RNA in the chorionic membrane at 7 dpi. (B) Corresponding H&E image. (C) IHC staining for cytokeratin highlights the trophoblast layer of the chorionic membrane showing that the ZIKV infection seen in (A) is in the trophoblasts of the chorionic membrane. Scale bars represent 100 µm.

**Fig 5.**
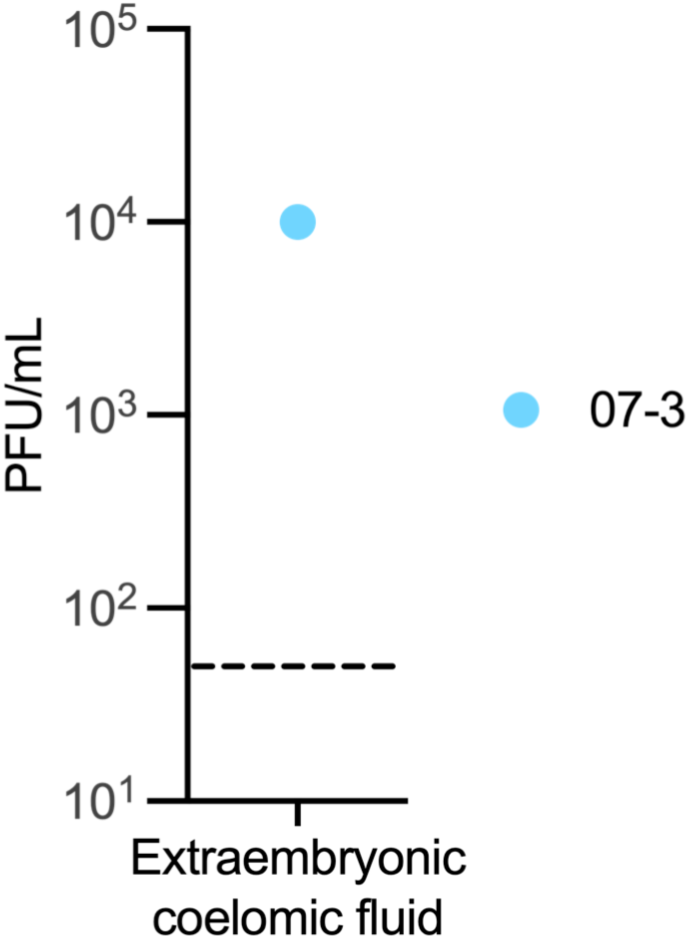
Infectious ZIKV in Extraembryonic coelomic fluid at 7 dpi. Infectious virus detected by plaque assay. The LOD of 50 plaque-forming units (PFUs) is represented by a dashed line.

### ZIKV infection spreads in the chorionic membrane and infects the mesenchymal core of the placental villi but not the STBs or CTBs at 14 dpi

At 14 dpi, all cases had ZIKV RNA detected in the decidua and the chorionic membrane (Fig 6). ISH results show extensive infection of the chorionic membrane mainly in the mesenchymal layer, with sparse infection in the trophoblasts of the chorionic membrane (Fig 7A–F). immunofluorescent staining (IF) revealed that a some of the positive cells were CD163-expressing macrophages, but most ZIKV signal is not found within these cells (Fig 7G, H). To determine if the ZIKV RNA detected by ISH represents replicating virus, we performed multiplex fluorescence *in situ* hybridization (mFISH) and found replicative intermediate RNA, suggesting that the ZIKV found in the chorionic membrane is replicating (S5 Fig). In addition to the infection of the chorionic membrane, infectious ZIKV RNA was detected in the extraembryonic coelomic fluid in the three cases, 14-1, 14-2, and 14-4 (Fig 8), 14-3 had ZIKV RNA detected in extraembryonic coelomic fluid (S4C Fig), but there was not enough of the fluid collected for a plaque assay.

**Fig 6.**
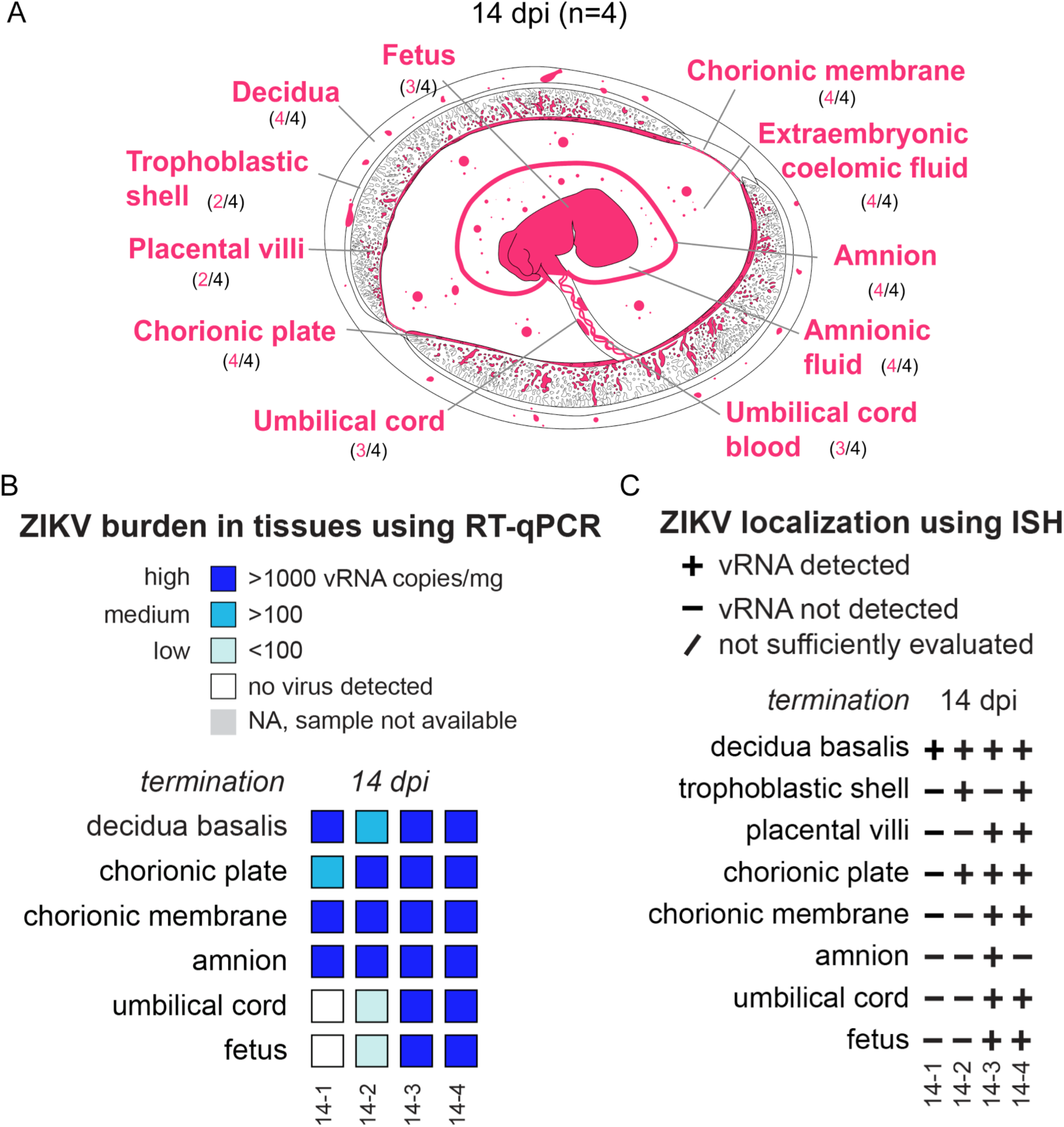
Summary of ZIKV viral burden at 14 dpi. (A) Artistic representation of ZIKV infection in the maternal-fetal interface at 14 days post infection (dpi). Pink tissue/structure names denote that ZIKV RNA was detected via ISH or RT-qPCR. The number of cases where the tissue had ZIKV RNA detected out of the number of cases in which tissue was evaluated is indicated below each tissue name. An artistic representation of the infection of each tissue is illustrated in pink in the figure. (B) Summary of the ZIKV RNA detected via RT-qPCR. The level of ZIKV burden is summarized as high (>1000 ZIKV RNA copies/mg of tissue, medium (>100 ZIKV RNA copies/mg or mL), low (<100 copies/mg of tissue), below the limit of detection, or NA, sample not available. The theoretical limit of detection for all the tissues is 3 copies/mg. (D) Summary of ZIKV RNA detected via ISH. ZIKV RNA was noted as present (+) or absent (-) in the indicated tissues/structures. In some instances, adequate histological samples were not obtained, and could not be properly evaluated (/). (B) (C) Individual animal IDs are listed on the bottom.

**Fig 7.**
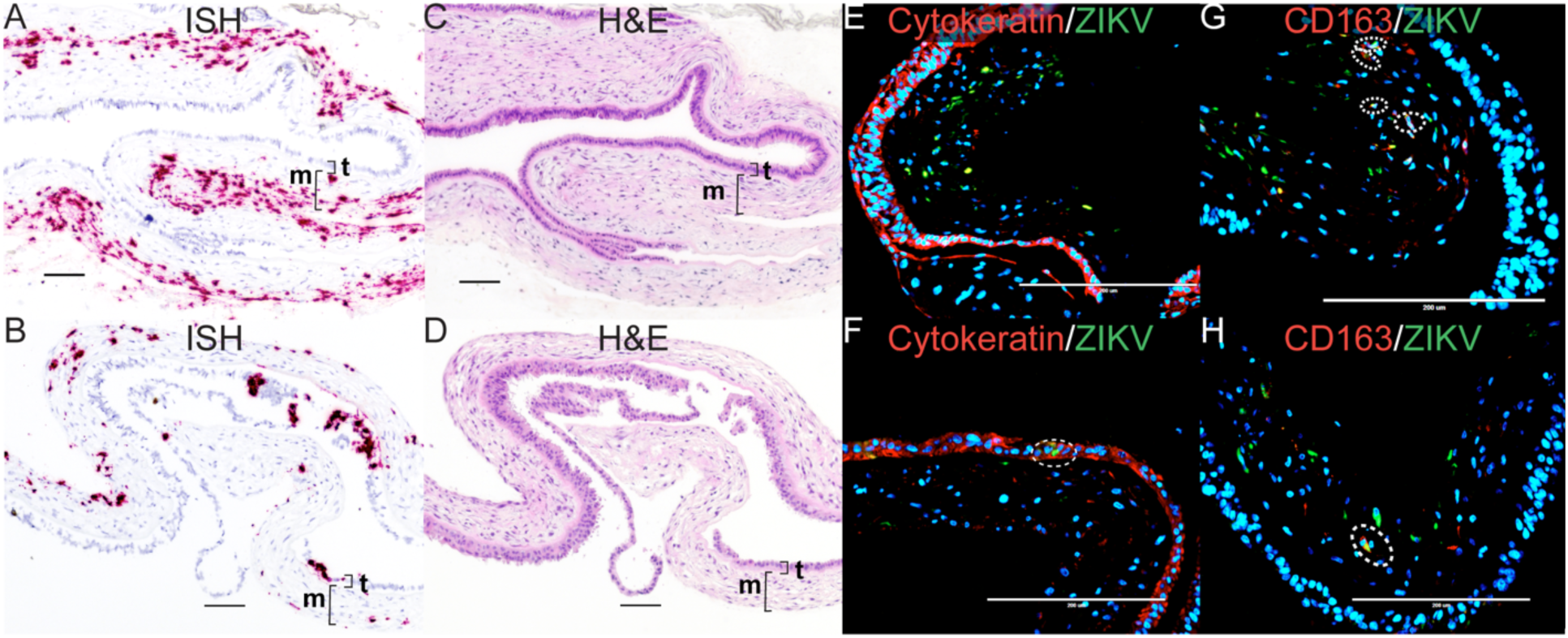
ZIKV infection in the chorionic membrane at 14 dpi. (A)(C)(E)(G) Images from 14-3 chorionic membrane. (B)(D)(F)(H) Images from 14-4 chorionic membrane. (A)(B) Pink staining showing ZIKV detected via ISH predominantly within the mesenchymal layer of the chorionic membrane. (C) Corresponding H&E stain of (A). (D) Corresponding H&E stain of (B). The mesenchymal layer is indicated by m and the trophoblast layer is indicated by t. (E)(F) IF staining for cytokeratin (red) and ZIKV (green), and DAPI nuclear stain (blue). (F) The dashed circle indicates the area of colocalization showing ZIKV infection in the trophoblasts of the chorionic membrane. (G)(H) IF staining with CD163 (red), ZIKV (green), and DAPI nuclear staining (blue). The dashed circles indicate areas of colocalization showing ZIKV infection in macrophages. (A)–(D)The scale bar represents 100 µm, (E)–(H) 200 µm.

**Fig 8.**
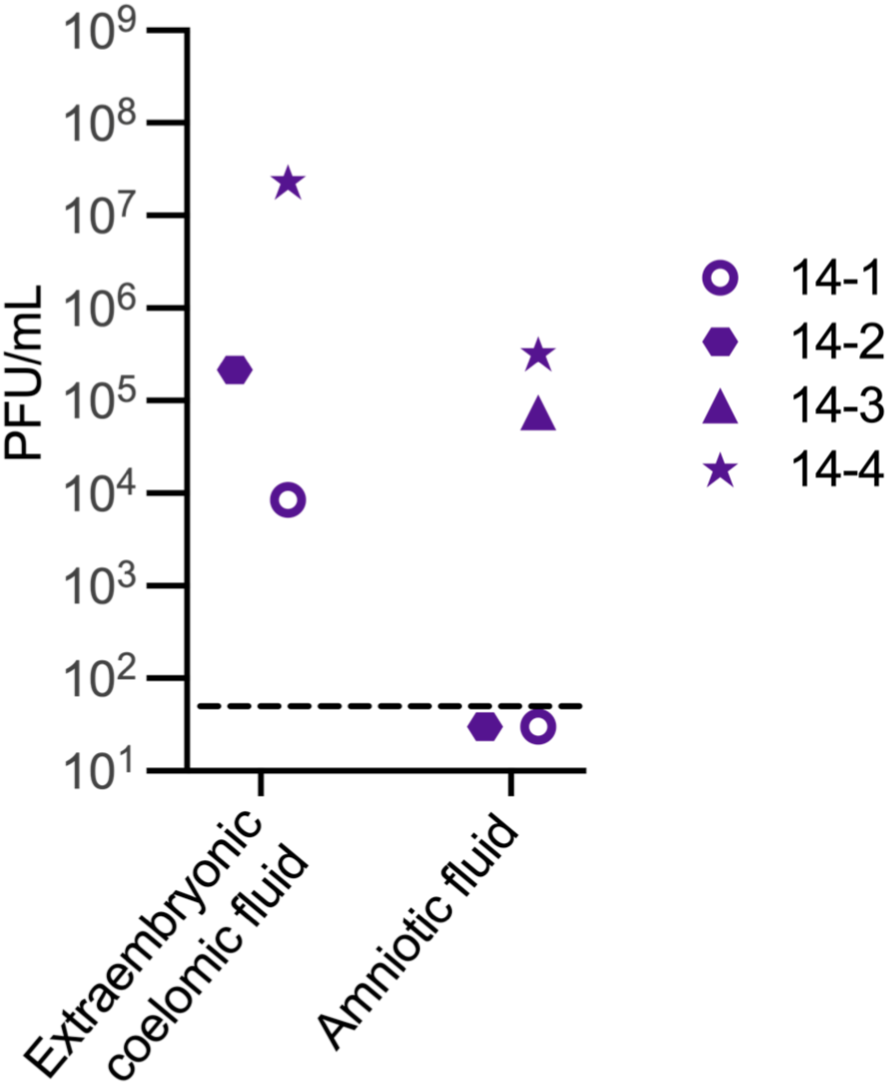
Infectious ZIKV in extraembryonic coelomic and amniotic fluid at 14 dpi. Infectious virus detected by plaque assay. The LOD of 50 plaque-forming units (PFUs) is represented by the dashed line.

At 14 dpi, ZIKV was detected in the chorionic plate in all four cases (Fig 6). ISH results show ZIKV RNA within the mesenchymal tissue of the chorionic plate, and a few of the positive cells were CD163-expressing macrophages (S6 Fig). ZIKV RNA was detected in the amnion in all 14 dpi cases (Fig 6). The cellular location of ZIKV in the amnion was challenging to determine due to the fragile nature of the membrane. The antigen retrieval process necessary for ISH and IF distorts the morphology of the cells. IF staining identified that some, but not the majority, of the ZIKV-infected cells in the amnion are CD163-expressing macrophages (S7 Fig). The amniotic fluid contained infectious ZIKV in two cases, 14-4 and 14-3, (Fig 8) and 14-2 had ZIKV RNA, but not infectious virus (S4D Fig). Infectious ZIKV in the amniotic fluid can easily infect the fetus in early gestation because it is absorbed by the fetus prior to skin keratinization [34].

ZIKV RNA was detected in the placental villi in the two cases where infectious ZIKV was present in the amniotic fluid (Fig 6). Importantly, ZIKV RNA was not detected in STBs or CTBs in the placental villi but was restricted to the mesenchymal core of the villi, mainly in the placental macrophages, Hofbauer cells (HBCs), where the virus was replicating (Fig 9).

**Fig 9.**
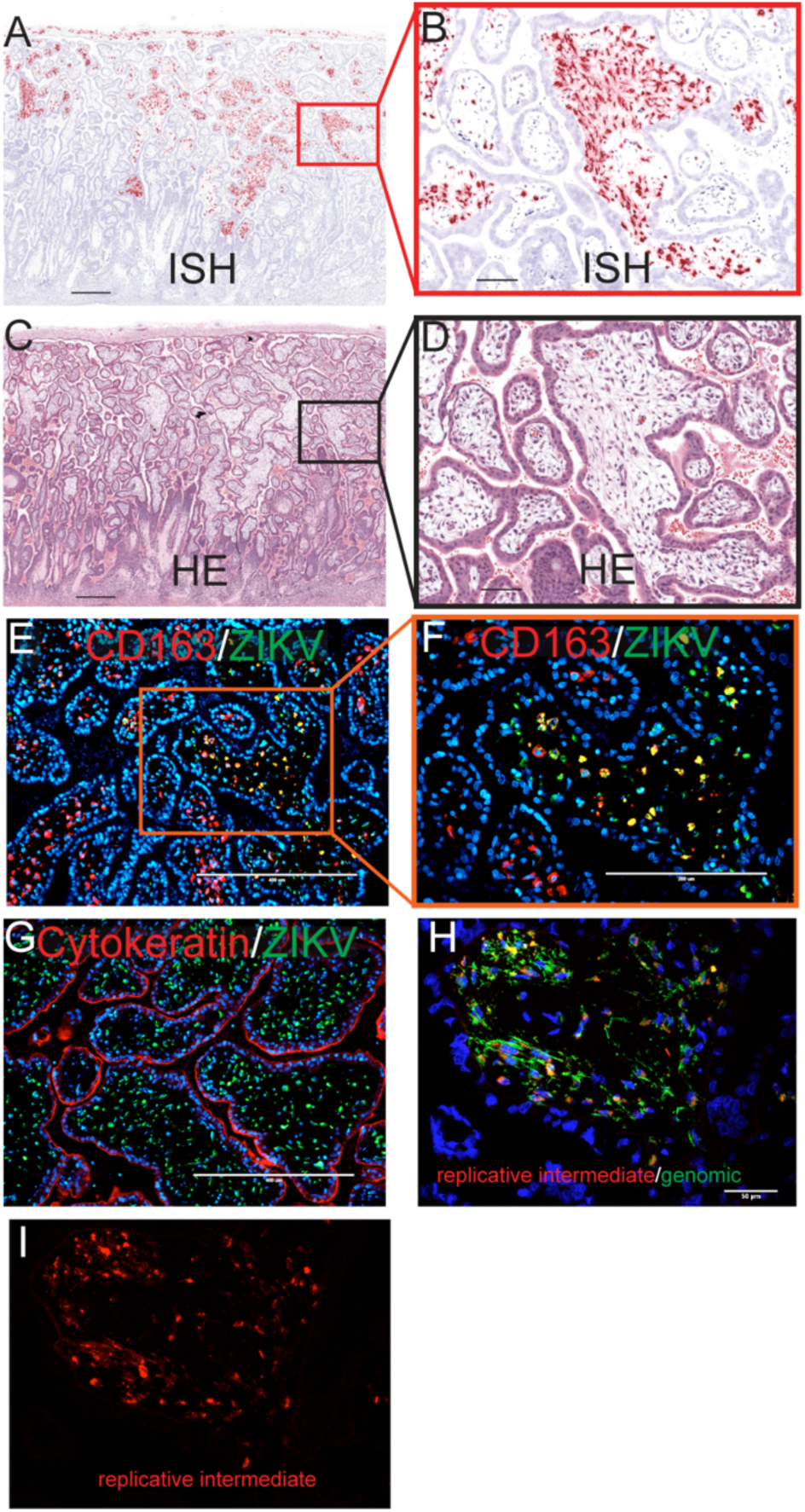
Infection in the placental villi at 14 dpi. (A) Pink staining shows ZIKV detected via ISH. (C) corresponding H&E section. The red square in (A) identifies the area magnified in (B) as does the black square in (C) for (D). (A)(C) The scale bars represent 500 µm. (B)(D) The scale bars represent 100 µm. (E) IF staining for CD163 (red) ZIKV (green) and DAPI (blue). The scale bar represents 400 µm. The orange square shows the portion that is magnificent in (F). (G) IF staining for cytokeratin (red) ZIKV (green) and DAPI (blue). (F)(H) The scale bars represent 400 µm. (H) Multiplex fluorescence in situ hybridization (mFISH) to detect genomic, positive sense ZIKV RNA (green) and replicative intermediate, negative sense RNA (red) with nuclear DAPI staining (blue). The scale bar represents 50 µm. The scale bar represents 100 µm. The replicative intermediate negative sense RNA (red) is shown alone in (J).

### ZIKV-DAK causes widespread infection of the fetus and fetal demise by 14 dpi

At 14 dpi, the two cases with infectious virus in the amniotic fluid, 14-4 and 14-3, had widespread fetal infection with ZIKV RNA detected throughout the fetus (Fig 10) as well as replicative viral intermediates (S8). One of the fetuses with widespread infection, 14-4, died *in utero* prior to surgical termination. The fetus was last confirmed to be alive four days prior to the day of fetectomy. Histopathological evaluation revealed fetal tissue autolysis. Due to the extent of viral infection in fetus 14-4, we believe ZIKV infection was the cause of the intrauterine demise. One case at 14 dpi, 14-2, had relatively low levels of ZIKV RNA detected in the fetal cerebral spinal fluid and a liver biopsy, whereas 14-1 did not have any ZIKV detected in the fetal tissues, fetal fluids, or the umbilical cord (Fig 6, Fig 10A, B). Interestingly, ZIKV RNA was only seen in the placental villi in cases with widespread fetal infection.

**Fig 10.**
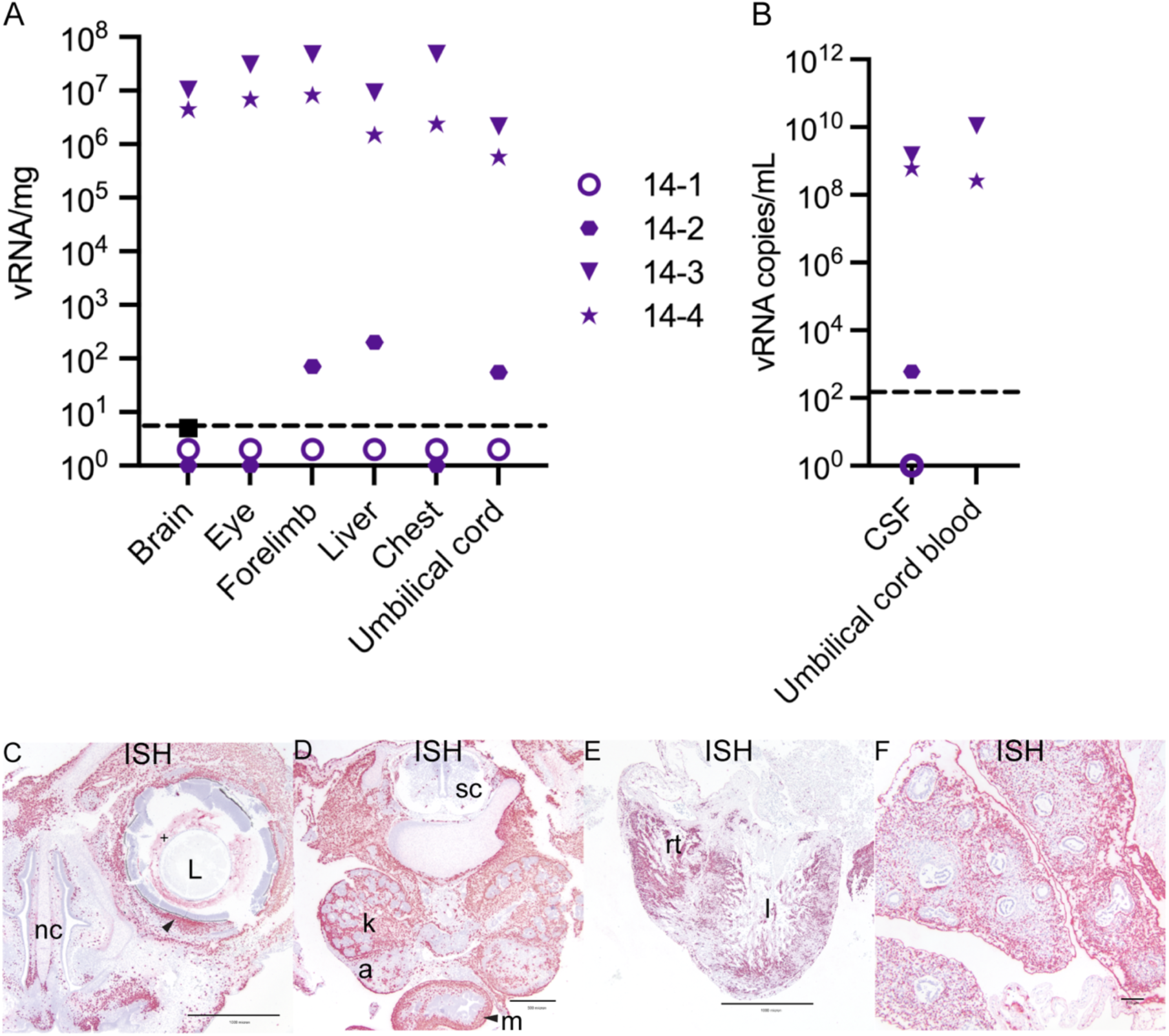
ZIKV burden in the fetus at 14 dpi. (A) ZIKV RNA detected in fetal tissues and organs. The limit of detection of 3 copies/mg is depicted by the dashed line. (B) ZIKV RNA detected in fetal fluids. The limit of detection of 150 copies/mL is depicted by the dashed line. CSF = cerebrospinal fluid. (C)–(F) ZIKV RNA detected via ISH (pink). (C)(D) images from 14-3 and (E)(F) from 14-4. (C) Infection detected in the eye of the fetus with ZIKV (+) in the material (vitreous/aqueous humor) surrounding the lens (L) and within the sclera of the eye indicated by the arrow. ZIKV was also detected subjacent to the nasal mucosa of the fetal nasal cavity (nc). Scale bar represents 1000 µm. (D) ZIKV RNA detected in the kidneys (k), adrenal glands (a), and intestinal muscular layers (m) with minimal scattered positivity in the spinal cord (sc). Scale bar represents 500 µm. (E) Abundant ZIKV was detected throughout the myocardium of the right (rt) and left (l) ventricles of the heart and (F) in the pulmonary interstitium of the lungs with sparing of the respiratory epithelium of the bronchi and bronchioles. (E) Scale bar represents 1000 µm and (F) 100 µm.

### ZIKV vertical transmission was further investigated in an additional cohort of pregnant dams

The fetectomy cohort evaluated the MFI and fetus at a relatively early stage of vertical transmission, at 7 dpi, when ZIKV had reached the MFI but not the fetus, and at a late stage, at 14 dpi, when ZIKV had already reached the fetus. To further map the emergence and distribution of ZIKV in the MFI tissues, pregnancies were terminated at 3, 6, 9, and 10 dpi. ZIKV inoculation of these dams was identical to the first cohort. In contrast to the first cohort, at the time of pregnancy termination the dams were anesthetized and perfused with 4% paraformaldehyde (PFA) through the heart. Once all tissues were fixed in situ, the intact gravid uterus was collected and further post-fixed.

The entire uterus was serially sectioned into coronal tissue slices using a standardized jig (slicing box). These coronal uterine sections were routinely processed and embedded in paraffin into extra-large histology cassettes. Cellular localization of ZIKV RNA was then evaluated using ISH and IF.

### ZIKV infects endovascular EVTs in the decidua by 6 dpi

ZIKV RNA was not detected at 3 dpi in the MFI or fetus (S9 Fig), however ZIKV RNA was detected at the 6 dpi, where ZIKV was found exclusively in the decidua (Fig 11). At 6 dpi, ZIKV was detected mainly within large decidual vessels (Fig 11A–C) resembling what was seen at 7 dpi. Immunofluorescence (IF) staining revealed that ZIKV was primarily localized to the endovascular EVTs (Fig 11D) and was found to be replicating (Fig 11E). ZIKV RNA was also located in non-trophoblastic cells in other parts of the decidua at 6 dpi (S10 Fig), but the infection in these areas appears less extensive than what was noted in the decidual vessels. The combined evaluation of tissues from the fetectomy and perfusion cohorts at 7 and 6 dpi suggests that ZIKV first infects the MFI by spreading from the maternal blood to the decidua.

**Fig 11.**
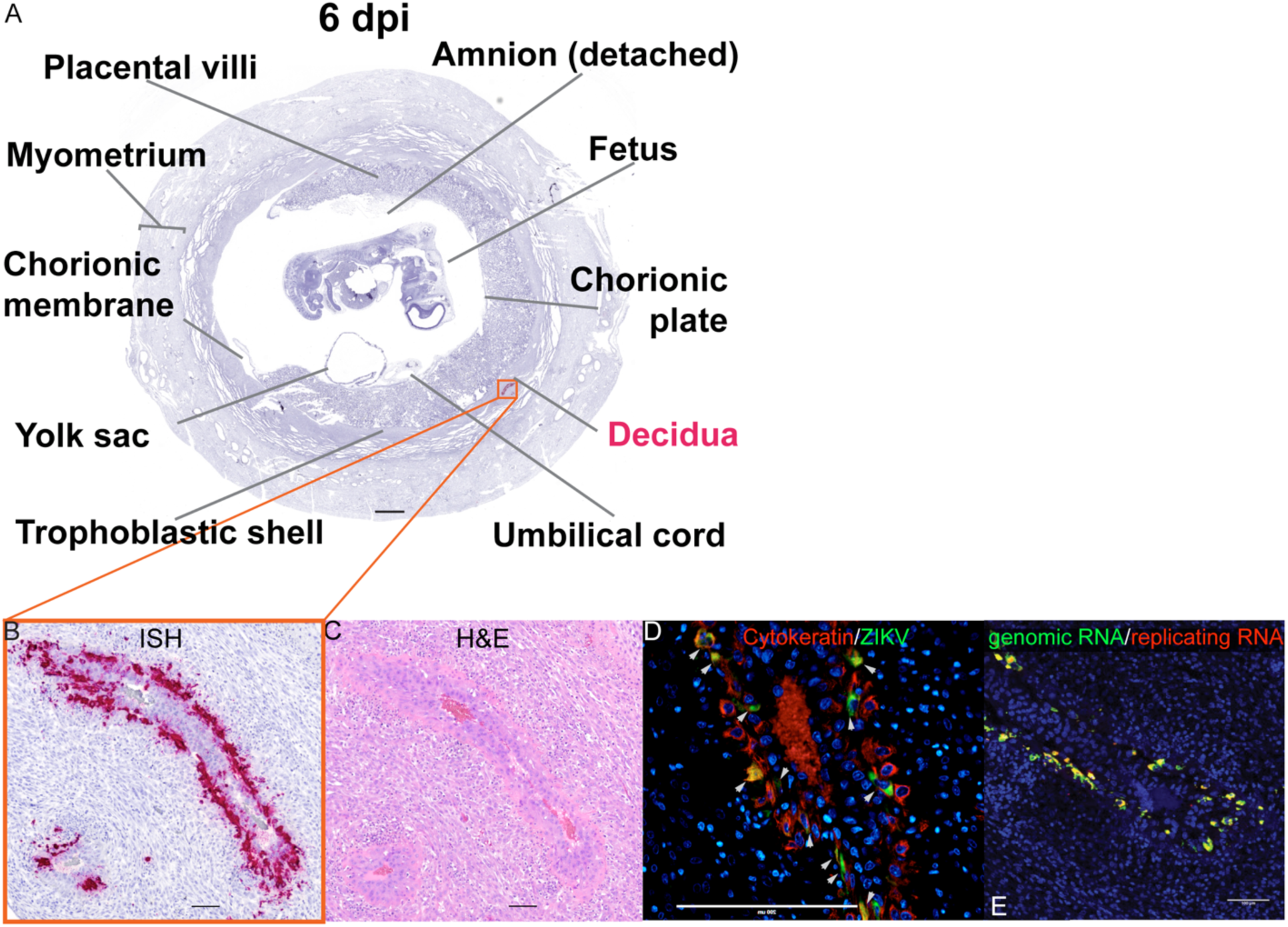
ZIKV infection in large decidual vessels at 6 dpi. (A) Photomicrograph of a representative coronal section of the gravid uterus evaluated via ISH for ZIKV RNA (pink). Pink tissue/structure names indicate the detection of ZIKV RNA via ISH. (B) Magnification of ZIKV RNA detected in a decidual vessel. (C) H&E staining of a serial section. (B)(C) The scale bar represents 100 µm. (D) IF staining of the same decidual vessel for cytokeratin (red), ZIKV (green), and DAPI for nuclear staining (blue). The scale bar represents 200 µm. (E) Multiplex fluorescence in situ hybridization (mFISH) to detect genomic, positive sense ZIKV RNA (green) and replicative intermediate negative sense RNA (red) with nuclear DAPI staining (blue). Colocalization of genomic RNA and replicating RNA (yellow) showing that the ZIKV RNA detected in the EVTs represents replicating virus. The scale bar represents 100 µm.

### ZIKV preferentially infects the trophoblasts in the chorionic membrane and at the peripheral margin of the placental disc

At 9 dpi, we detected relatively sparse ZIKV RNA in the decidua, the chorionic membrane, and the peripheral margin of the placental disc (Fig 12). In the chorionic membrane, we see ZIKV RNA in the trophoblast layer (Fig 12A, B). Adjacent to ZIKV positive trophoblasts of the chorionic membrane, positive staining was observed at the peripheral margin of the placenta, the border where the placenta ends and the chorionic membrane starts (Fig 12B). Evaluation of this area of the placenta was limited in the fetectomy cohort as the architecture was disrupted by the sampling process. The infection in the peripheral margin of the placenta and the chorionic membrane was only seen in the slice of the uterus shown in Fig 12. In the decidua, we observed relatively sparse ZIKV infection at 9 dpi (Fig 12C) compared to 6 and 7 dpi.

**Fig 12.**
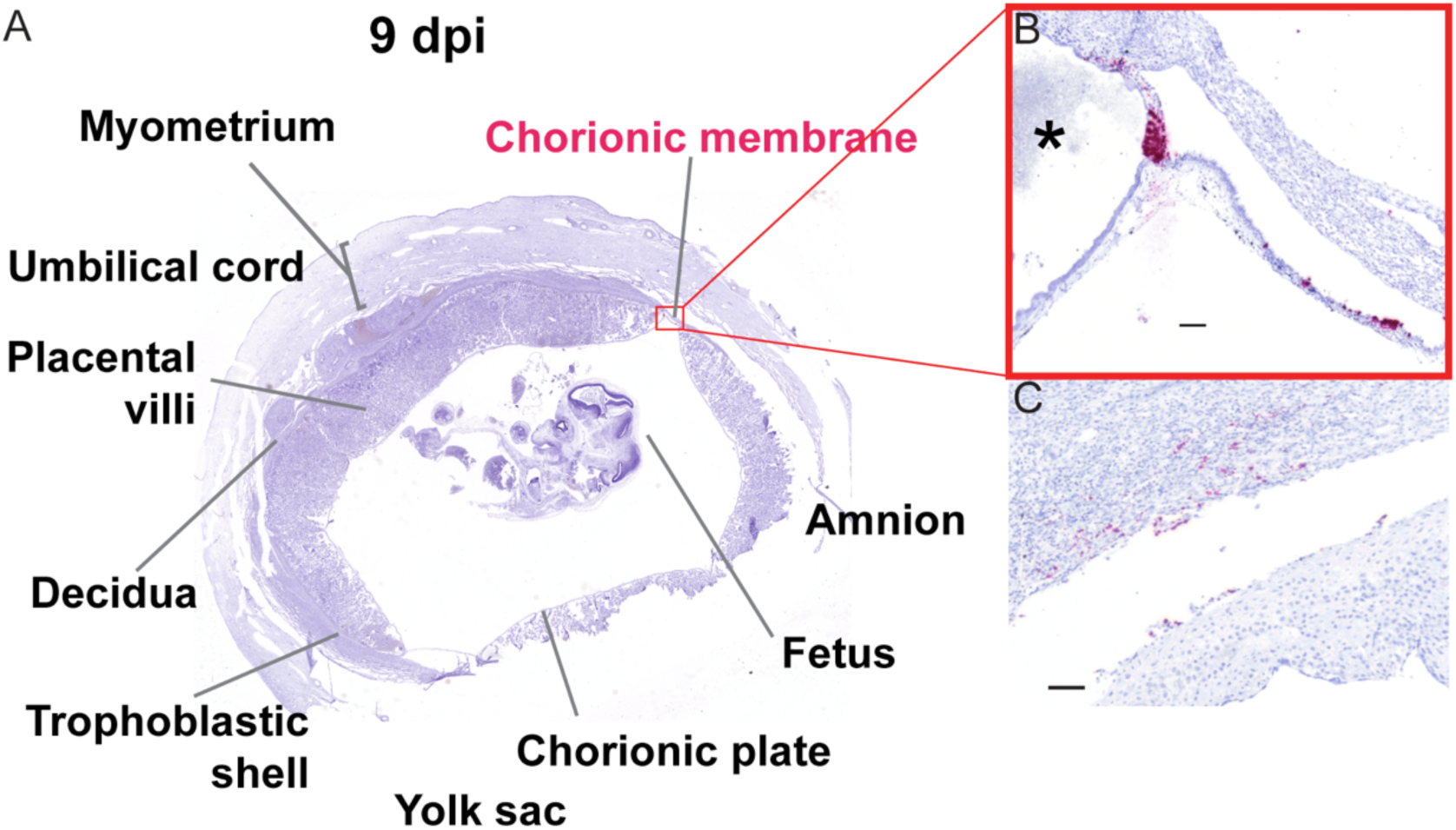
ZIKV infection of chorionic membrane and peripheral margin of the placenta at 9 dpi. (A) Photomicrograph of a representative coronal section of gravid uterus evaluated via ISH for ZIKV RNA (pink). Pink tissue/structure names indicate the detection of ZIKV RNA via ISH. (B) Magnification of ISH staining showing ZIKV RNA in the trophoblasts of the chorionic membrane and in the peripheral margin of the placenta. * indicates the placenta. (C) Sparse infection of the decidua. (B)(C) The scale bar represents 100 µm.

In the pregnancy collected at 10 dpi, the MFI and fetus were highly infected (Fig 13). Infection was again observed in decidual vessels (S11A–C Fig). Evaluation of the intact gravid uterus allowed for unique observations that were not possible in the fetectomy cohort; for example, coronal slices of the superior pole of the uterus highlights the extent of ZIKV infection in the EVTs of the trophoblastic shell (S12 Fig). At 10 dpi, ZIKV RNA was detected again in the chorionic membrane (Fig 13A–E). IF staining with cytokeratin emphasizes that ZIKV infection at 10 dpi is restricted to the trophoblasts in the chorionic membrane, with little to no ZIKV detected in the mesenchymal layer (Fig 13B, C). Compared to the pattern of infection observed at earlier time points, by 10 dpi the infection of the chorionic membrane appears to be more expansive (Fig 13D, E). ZIKV RNA was detected in the chorionic membrane in all slices of that uterus that contained the membrane. ZIKV was also seen in the trophoblasts at the peripheral margin of the placenta in the most exterior trophoblasts adjacent to the decidua (Fig 13F, G). This pattern of infection was seen in multiple slices of the uterus.

**Fig 13.**
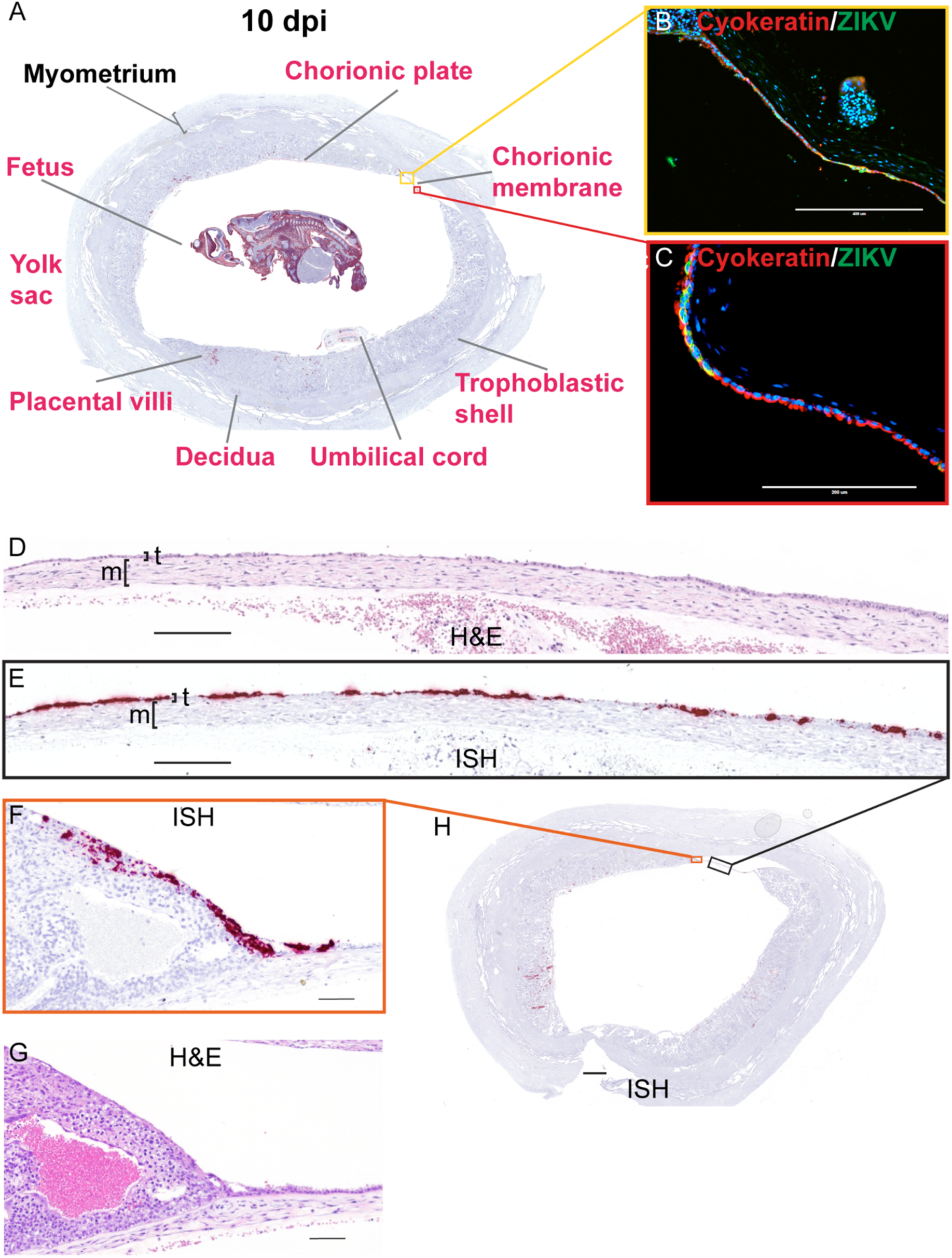
ZIKV infection at 10 dpi shows extensive infection of the chorionic membrane. (A) A Representative coronal section of gravid uterus evaluated via ISH for ZIKV RNA (pink). Pink tissue/structure names indicate the detection of ZIKV RNA via ISH. The fetal head had been added to this image from a different coronal section. The yellow and red squares outline the areas magnified in (B) and (C), respectively. (B)(C) IF staining for cytokeratin (red) and ZIKV (green) and nuclear DAPI (blue). Colocalization of green and red shows that ZIKV has infected the trophoblasts of the chorionic membrane. (D) H&E stained section corresponding to section shown in (E) depicting ZIKV RNA detected via ISH (pink) in the trophoblast layer (t) of the chorionic membrane. The mesenchymal layer is indicated by m. (F) ZIKV RNA detected via ISH in the peripheral margin of the placental disc. (G) H&E stained section. (H) Full slide evaluated via ISH the black square outlines the portion magnified in (E) and the orange square outlines the portion magnified in (F). The scale bar represents 400 µm in (B), 200 µm in (C)(D)(E), 100 µm in (F)(G), and 2350 µm in (H).

### The presence of ZIKV at 10 dpi further supports a paraplacental route of vertical transmission

ZIKV is found in the villous core at 10 dpi, similar to the 14 dpi results, where the infection is limited to the mesenchymal tissue and not seen in either the STBs or CTBs of the placental villi (Fig 14 A–D). IHC and IF show that some of the infection is in HBCs, but a substantial amount of the infection looks to be outside HBCs in the stromal tissue of the villi (Fig 14C–F) and some infection of endothelial cells in the fetal vessels of the placental villi (Fig 14C, D). ZIKV is also found in the mesenchymal tissue of the chorionic plate (Fig 14B). IF staining shows that most ZIKV-positive cells in the chorionic plate are macrophages (Fig 14G), however we do see some infection in the endothelial cells of the fetal vessels (S13 Fig). The detection of replicative intermediate ZIKV RNA in the chorionic plate and placental villi suggests active viral replication in these sites (S14A–C Fig). Although ZIKV was detected in the trophoblasts of the chorionic membrane, and trophoblasts on the peripheral margin of the placental discs, ZIKV was not commonly seen in the adjacent trophoblasts of the chorionic plate. The chorionic plate’s trophoblasts were infected in only one coronal uterine slice taken from 10-1(S15 Fig).

**Fig 14.**
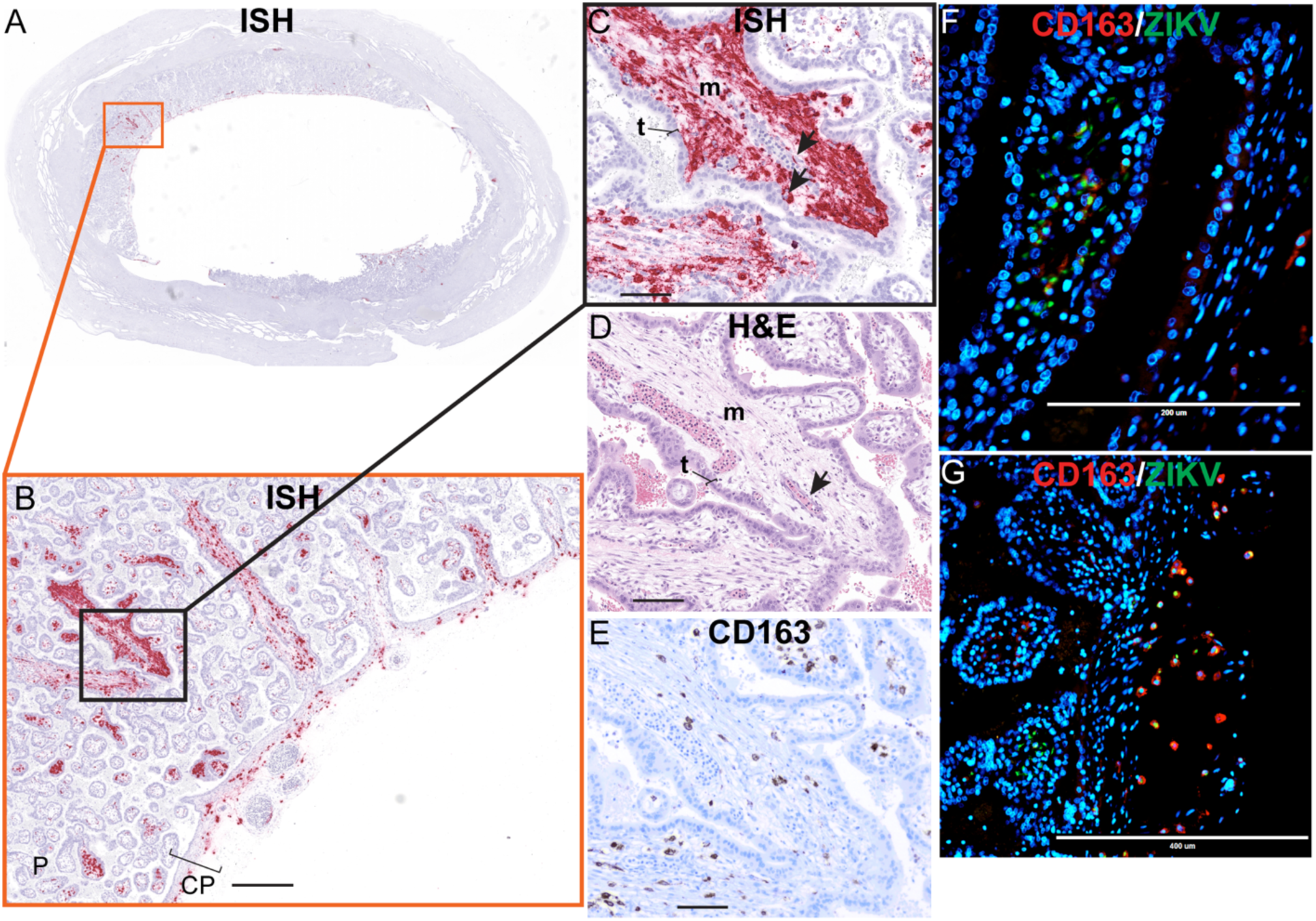
Mesenchymal tissue of the chorionic plate and villi infected with ZIKV at 10 dpi. (A) full slide evaluated with ISH shown as pink staining. The orange square outlines the area of the slide magnified in portion (B). (B) CP indicated chorionic plate. The black square outlines the portion of the slide magnified in (C-E). (D) H&E of the corresponding section is shown in (C). (C)(D) Arrows indicate endothelial cells, m indicates mesenchymal tissue and t indicates trophoblasts in the placenta. (E) Corresponding section with IHC staining for CD163. (F)(G) IF staining for CD163 (red) ZIKV (green) and DAPI nuclear staining (blue). (G) Photomicrograph of the placental villi with colocalization of green and red showing that ZIKV is within Hofbauer cells in the placenta villi, but not exclusively. (F) Depicts the chorionic plate of the placenta where ZIKV almost exclusively co-localized with CD163 showing ZIKV predominantly within macrophages in the chorionic plate. Scale bars represent 500 µm in (B), 100 µm in (C)(D)(E), 400 µm in (F), and 200 µm in (G).

Unfortunately, the amnion is extremely thin and fragile at this early stage of gestation and was lost to evaluation in this case. Assessment of the intact gravid uterus did, however, allow us to evaluate the residual secondary yolk sac. ISH revealed extensive ZIKV infection of the yolk sac, mainly within the extraembryonic mesoderm layer and within sites of fetal primary hematopoiesis (Fig 15).

**Fig 15.**
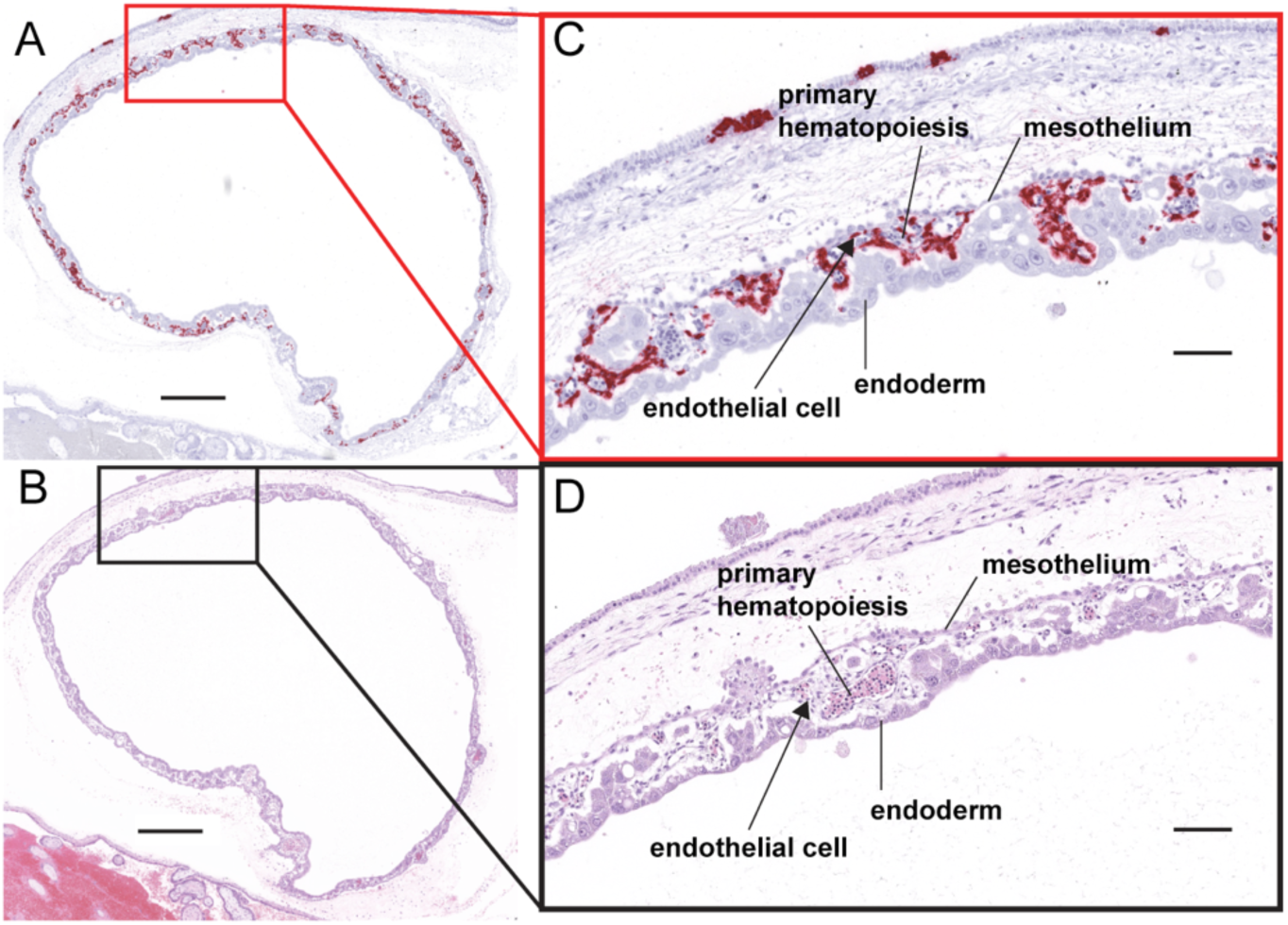
ZIKV infection in the yolk sac. (A) ISH staining for ZIKV shown as pink staining. Red square indicates the portion of the section magnified in (C). The scale bar represents 100 µm. (B) Corresponding H&E stained section. Black square indicates the portion of the section magnified in (D). (C)(D) Structures of the yolk sac outlined. The scale bars represent 500 µm in (A)(B) and 100 µm in (C)(D).

At 10 dpi, ZIKV was seen in the umbilical cord and fetus (Fig 16). ZIKV infection of the umbilical cord is primarily within the mesenchymal tissues (Fig 16A,B). The pattern of ZIKV infection in our perfused 10 dpi case, 10-1, reflects that seen in the two 14 dpi fetectomy cases with widespread infection. ISH evaluation of the intact fetus shows us the extent of fetal infection seen with ZIKV-DAK; the only structures in the fetus that were not infected appeared to be immature bone and hepatocytes in the liver (Fig 16C–J). ZIKV replicative intermediates were confirmed with mFISH (S14D, E Fig). The fetus was confirmed to have a heartbeat before the dam was euthanized, however infection would have likely compromised the fetus’s ability to progress to full term.

**Fig 16.**
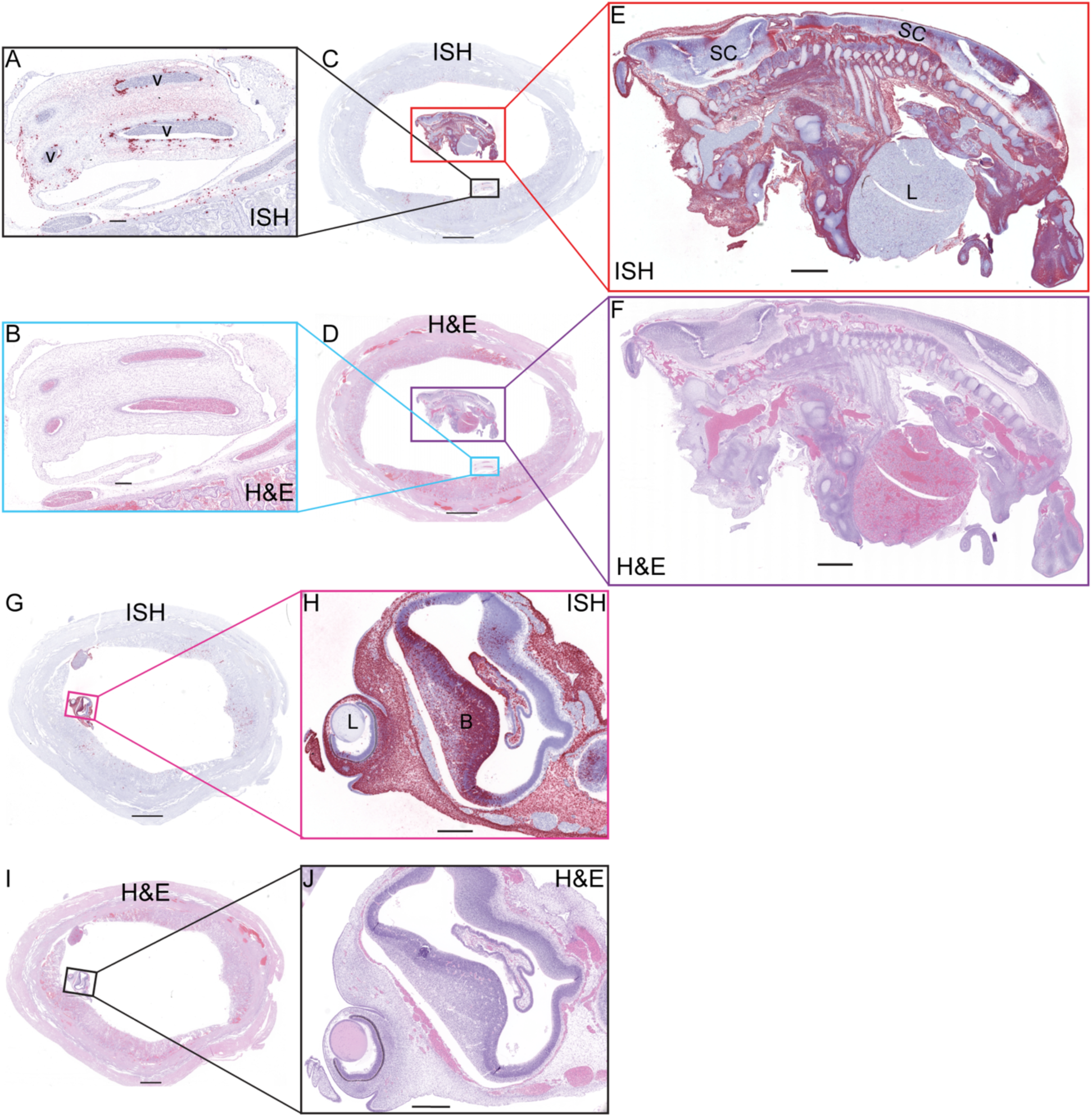
ZIKV RNA detected at 10 dpi via ISH. (A) ISH staining for ZIKV (pink) in the endothelium lining umbilical vessels (v) and multifocally scattered throughout Wharton’s jelly surrounding the affected vessels of the umbilical cord. (B) Corresponding H&E stained section. (C) Full slide evaluated for ZIKV RNA via ISH. The black square outlines the area magnified in (A) and the red square is magnified in (E). (D) The corresponding H&E stained section shown in (B). The area outlined in blue is magnified in (B) and the area outlined in purple is magnified in (F). (E) ZIKV RNA detected in the body of the fetus via ISH (pink). There is marked diffuse ZIKV RNA (pink) in the majority of tissues with less intense and single cell positivity in the spinal cord (SC) and liver (L). ZIKV RNA was not detected in the cartilage and immature bone of the skeleton of the fetus. (F) The corresponding H&E stained section of the sample shown in (E). (G) Full slide evaluated for ZIKV RNA via ISH. The pink square outlines the portion of the slide magnified in (H). (H) The head of the fetus with ZIKV RNA identified via ISH (pink) in the sclera of the eye (arrow), brain (B), and tissues of head. ZIKV RNA was not found in the lens (L) of the eye. (I) (J) Corresponding H&E stained section of specimens shown in (G) and (I), respectively. (A)(B) Scale bars represent 250 µm. (C)(D)(G) The scale bars represent 5000 µm. (E)(F) The scale bars represent 1000 µm. (H)(J) The scale bars represent 500 µm. (I) The scale bar represents 3450 µm.

### ZIKV-DAK infection and vertical transmission have minimal impact on placental function

Histological evaluation of the placenta was performed by a board-certified pathologist (TKM) who specializes in human placental pathology. To determine what effect ZIKV-DAK has on the placenta, we evaluated full thickness placental sections from the eight cases in the fetectomy cohort (7 and 14 dpi) compared to eight gestational age-matched controls. The histopathological evaluation revealed abnormalities in two placentas (pregnancy 14-2 and 14-4) collected at 14 dpi. There was a slight increase in acute infarctions in the placenta and early maternal decidual vasculitis, but these pathologies are also seen in control placentas (S1 Table). Only the case with *in utero* fetal demise, 14-4, had unique pathological findings absent from any of the controls; this case had remote infarctions in the placenta and multifocal maternal decidual vasculitis (S1 Table). To further evaluate the impact of ZIKV infection on the function of the placenta, we measured plasma levels of steroid pregnancy hormones for the eight dams from the first cohort and compared them to controls. Consistent with our histopathologic findings, ZIKV infection did not affect estradiol or progesterone levels (S16 Fig) suggesting there was not a significant impact on STB function.

### Increased IFNλ-1 mRNA expression in ZIKV placentas

Type III interferons expressed by STBs have been shown to protect the placental trophoblasts from ZIKV infection [16, 35]. In particular, IFNλ-1 expression has shown to protect primary human trophoblast cells and tissue explants from ZIKV infection [16, 35]. To better understand what role type III interferon expression in the placenta might play in protecting STBs from infection, we analyzed the expression of IFNλ-1 mRNA via RT-qPCR in placentas from the eight pregnancies in the fetectomy cohort and three control pregnancies (Table 1). IFNλ-1 mRNA expression was only detected in placenta samples from 07-3, 14-3, and 14-4 (Table 1). Interestingly, out of the four 14 dpi samples IFNλ-1 mRNA was only detected in the two cases that had infection in the placental villi. IFNλ-1 mRNA was also found in one 7 dpi case, 07-3, the same case that had infectious ZIKV in the extraembryonic coelomic fluid. Additionally, we analyzed IFNλ-1 mRNA expression in the remaining chorionic membrane RNA samples collected from six infected pregnancies. Interestingly, IFNλ-1 mRNA expression was only detected in the three 14 dpi samples (14-2, 14-3, 14-4) but was not detected in any of the 7 dpi samples (7-1, 7-2, 7-7) (Table 1).

**Table 1.**
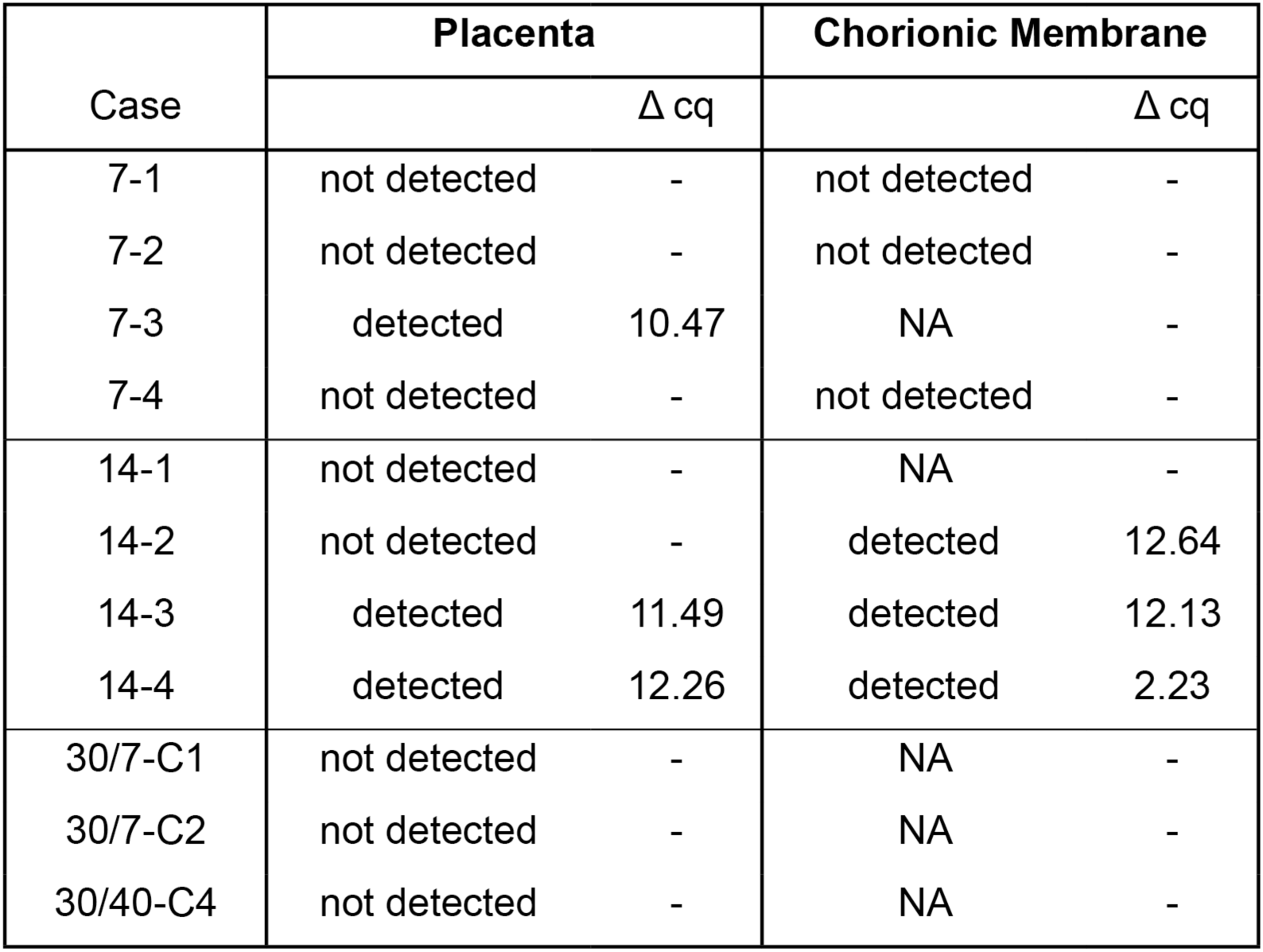
IFNλ-1 mRNA expression in placenta and chorionic membrane homogenates as detected by RT-qPCR. For samples with detectable IFNλ-1 mRNA the ΔCq. ΔCq = the cycle of quantification for IFNλ-1 -the cycle of quantification for the housekeeping gene beta-actin. NA = sample not available.

## Discussion

### ZIKV-DAK vertically transmits through the fetal membranes

In this study, we inoculated 12 ZIKV-naive pregnant rhesus macaques with 10^4^ PFU of ZIKV-DAK at approximately gd 30 during the late embryonic period of gestation. We then terminated the pregnancies at 3 to 14 dpi to provide snapshots at multiple time-points throughout the stages of infection in the MFI and fetus, allowing us to determine the order in which specific tissues and cells become infected as the virus passes from dam to fetus. The timing and pattern of ZIKV infection in the MFI and fetus suggests that ZIKV-DAK vertically transmits through the fetal membranes. ZIKV can reach the MFI by 6 dpi and infect the fetus by 10 dpi, although the precise timing of fetal infection varies between animals. We found that infection of the chorionic membrane and the extraembryonic coelomic fluid preceded infection of the fetus and the mesenchymal tissue of the placental villi. Importantly, we did not find any evidence to support a transplacental route of ZIKV vertical transmission in which ZIKV would reach the fetus by infecting the STBs and then the CTBs of the placental villi. ZIKV was not detected in STBs or CTBs in placental villi at any time. This study provides evidence to support that ZIKV-DAK is vertically transmitted through the chorionic membrane in rhesus macaques.

We uniquely used two complementary approaches to evaluate the conceptus post-ZIKV infection to better understand how ZIKV-DAK overcomes MFI barriers and reaches the fetus. The fetectomy cohort provided evaluation of the MFI at relatively early (7 dpi) and late (14 dpi) vertical transmission stages. RT-qPCR was used to broadly survey which tissues and fluids contained ZIKV RNA. ISH and IF were then used to determine what specific structures and cells were infected with ZIKV. Evaluation of additional time points with the perfusion cohort provided a histologically complete picture of ZIKV vertical throughout the entire MFI and conceptus within an individual pregnancy. This technique provided new insight into how ZIKV may disseminate within and across specific tissues, which areas are more vulnerable to ZIKV infection, and offers the opportunity to evaluate tissues that may not be collected during fetectomy.

ZIKV detection was limited to the mesenchymal core of the placental villi. ZIKV infection was only seen in the placental villous core in cases with overwhelming fetal infection. Additionally, there appeared to be more virus in the core of the villi closest to the fetal side of the placenta. These results may suggest that the mesenchymal core of the placenta is being infected through the fetal circulation. It is plausible that infection of the villous core is only secondary to fetal infection, and warrants further examination. Infection of the peripheral margin of the placental discs, observed in the terminal perfusion cohort in this study, may also present an additional pathway of ZIKV vertical transmission that has not been previously revealed by other studies. It is possible that the trophoblasts in the peripheral margin of the placenta are phenotypically similar to the trophoblasts in the chorionic membrane. The data presented in this study support the conclusion that ZIKV is vertically transmitted by infecting the trophoblasts in the chorionic membrane to reach the fetus, bypassing the more robust barrier of the placental STBs.

The susceptibility of the chorionic membrane trophoblasts, but not the STBs and CTBs of the placenta villi may be explained by differences in IFNλ-1 expression. Placental expression of IFNλ-1 mRNA suggests that in early gestation IFNλ-1 is not constitutively expressed or was weak and undetectable by RT-qPCR yet capable of increasing expression in response to viral infection. In the chorionic membrane, IFNλ-1 mRNA expression was detected in all three of the 14 dpi (gd 44) samples evaluated but not in any of the three 7 dpi (gd 37) samples. However, because we were unable to evaluate control samples, we do not know if upregulation occurred in response to infection or in response to normal developmental changes. This pattern of IFNλ-1 mRNA expression in the chorionic membrane may help explain why ZIKV appears to infect the chorionic membrane trophoblasts at 7 and 10 dpi, but at 14 dpi the infection is seen predominantly in the mesenchymal layer of the membrane. However, our evaluation of IFNλ-1 is extremely limited in this study, and further investigation is warranted. Previously published ZIKV studies found that inoculated pregnant rhesus macaques with the same dose of ZIKV-DAK resulted in a high rate of vertical transmission and fetal demise [28], but when inoculated at gd 45, pregnancies were viable until term and vertical transmission was not detected [23]. These findings indicate that there are gestational changes between gd 30 and gd 45 that dramatically decreases the vulnerability of the conceptus to ZIKV. These changes may be related to the susceptibility and permissiveness of chorionic membrane trophoblasts to ZIKV, but further investigation is needed.

### The use of African-lineage ZIKV

In this study, we used ZIKV-DAK, an African-lineage ZIKV isolate, to infect at gd 30 because these methods produce a high rate of vertical transmission within a discrete period of time [28]. The use of ZIKV-DAK may be a shortcoming of this model, because African-lineage ZIKV has not yet been reported to cause adverse pregnancy outcomes in humans. However, ZIKV-DAK effectively models ZIKV vertical transmission using a genetically diverse, immunologically competent, and physiologically relevant animal model [28,36,37]. Additionally, *ex vivo in vitro* studies suggest that the overall susceptibility of essential barrier tissues (STBs in placenta villi, and the chorionic membrane) is not different between Asian and African ZIKV isolates [10,16,17,20], although with *in vitro* and *ex vivo* studies, the cytopathic effect or the viral titers reached may be different between African versus Asian/American-lineage ZIKV [17, 20].

Importantly, all studies that used ZIKV of multiple lineages found that if one lineage of ZIKV was able to infect STBs or chorionic membrane, the other was able to as well [10,16,17,20].

### Infection of intravascular EVTs and yolk sac may have important implications for fetal outcomes

In contrast to Asian/American lineage ZIKV, vertical transmission of ZIKV-DAK is more likely to cause fetal demise rather than congenital defects [28,38,39]. The overwhelming infection seen in the fetuses in this study and others [14, 28] supports the conclusion that fetal demise occurs from fetal ZIKV infection.

To our knowledge, this is the first study to report productive infection of the yolk sac by ZIKV. We observed that ZIKV is able to infect the yolk sac, particularly within the sites of fetal primary hematopoiesis that give rise to immune cell populations including microglia [40]. ZIKV infection of the yolk sac could impact normal fetal immune development and may facilitate viral dissemination to the fetus.

Although STBs appear to be resistant to ZIKV infection, endovascular EVTs appear to be particularly vulnerable to ZIKV infection. In this study, we found large decidual vessels occluded by EVTs that were highly infected with ZIKV. If ZIKV infection kills or impairs intravascular EVTs, this would likely impact spiral artery remodeling [41]. Spiral artery remodeling is essential to a successful pregnancy; insufficient spiral artery remodeling has been linked to adverse pregnancy outcomes such as fetal growth restriction which has been reported in clinical cases of ZIKV infection during pregnancy [9]. We do not know why EVTs are susceptible to ZIKV infection while STBs are not. This difference in susceptibility may reflect differences in innate immune protection, namely type III interferon responses. Alternatively, it may reflect a difference in the ability of ZIKV to attach and enter these different trophoblasts. STBs also have a brush border of a dense layer of cortical actin, making pathogen attachment and entry difficult [42]. EVTs and STBs also express different surface antigens. For example, EVTs, but not STB, express CD56, a known ZIKV receptor [43]. Further studies are needed to understand what makes specific trophoblast populations (endovascular EVTs, EVTs in the trophoblastic shell, and trophoblasts in the chorionic membrane) vulnerable to ZIKV infection, while STBs and villous CTBs are not. Currently, we have limited knowledge regarding the barrier functions of these different trophoblast populations throughout gestation. Understanding these characteristics is essential to our knowledge of how vertical transmission of ZIKV and other pathogens occurs.

### Chorionic trophoblasts are a critical barrier to pathogens

The pathway of vertical transmission for many antenatal vertically transmitted TORCH pathogens is unknown [42, 44]. Colloquially, the transmission of many TORCH pathogens is referred to as transplacental, even though few pathogens have been shown to infect STBs [42,44,45]. STBs have been shown to release type III IFNs that are capable of protecting not only STBs but also neighboring cells from infection with ZIKV, rubella, cytomegalovirus (CMV), varicella zoster virus, and herpesvirus (HSV-1) [16,35,46,47]. Some pathogens, however, have been shown to infect STBs. Hepatitis E genotype 1 (HEV-1), for example, can directly infect STBs by antagonizing type III IFN production [48].

When studying antenatal vertical transmission and congenital infection, the fetal membranes are often overlooked [12,49–52]. Similarly, our knowledge of the chorionic membrane trophoblasts lags considerably behind our understanding of the STBs and CTBs in the placental villi. In particular, we know little about their function as protective barriers against pathogens despite the fact that these trophoblasts line the majority of the conceptus in human pregnancy. Marsh et al 2022 evaluated trophoblasts in the chorionic membrane from second trimester human pregnancies and found that these trophoblasts in particular expressed *IFITM2*, a restriction factor preventing entry of viruses into cells [53]. However, a critical knowledge gap still remains. Improving modeling to evaluate the potential of current and future pathogens to include both transplacental and paraplacental antenatal vertical transmission pathways is essential. Our findings in this study demonstrate that the fetal membranes are a critical and vulnerable barrier to antenatal vertical transmission that warrants further investigation.

## Methods

### Experimental Design

Twelve pregnant female rhesus macaques were subcutaneously inoculated with 10^4^ PFU of a Senegal isolate of African-lineage Zika virus ZIKV/*Aedes africanus*/SEN/DAK-AR-41524/1984 (ZIKV-DAK), Genbank accession number KY348860, at approximately gestational day (gd) 30. Eights of these dams were included in the fetectomy cohort: had their pregnancies surgically terminated seven days post infection dpi, and four dams had their pregnancies surgically terminated at 14 dpi. There were also four dams in the perfusion cohort that received terminal perfusions with 4% PFA at 3, 6, 9 and 10 dpi. A total of eight pregnant dams were inoculated with saline to serve as controls. Four of these control pregnancies were terminated at approximately gd 37 (seven days post saline inoculation), and four were terminated at approximately gd 44 (14 days post saline inoculation). Dams with control pregnancies were subjected to the same experimental sampling regimen as the ZIKV infected pregnancies. Details on the timing of ZIKV/saline inoculation and pregnancy termination/necropsy are provided in S1 Table.

There was one additional dam that was inoculated with ZIKV-DAK that was excluded from the study. This dam, 14-5, never developed plasma viremia above the limit of detection (S17A Fig). We found that this dam had nAb levels prior to inoculation that was minimally expanded at 27 dpi (S17B, C Fig), suggesting that this dam was not immunologically ZIKV naïve and was not infected by the ZIKV-DAK inoculation. Therefore, we excluded this dam from the study. To avoid confusion, we have not assigned this dam to either cohort and do not discuss this dam outside of the methods.

### Ethics

The rhesus macaques used in this study were cared for by the staff at the Wisconsin National Primate Research Center (WNPRC) according to regulations and guidelines of the University of Wisconsin Institutional Animal Care and Use Committee, which approved this study protocol (G005691) in accordance with recommendations of the Weatherall report and according to the principles described in the National Research Council’s Guide for the Care and Use of Laboratory Animals. All animals were housed in enclosures with at least 4.3, 6.0, or 8.0 sq. ft. of floor space, measuring 30, 32, or 36 inches high, and containing a tubular PVC or stainless steel perch. Each individual enclosure was equipped with a horizontal or vertical sliding door, an automatic water lixit, and a stainless steel feed hopper. All animals were fed using a nutritional plan based on recommendations published by the National Research Council. Twice daily, macaques were fed a fixed formula of extruded dry diet (2050 Teklad Global 20% Protein Primate Diet) with adequate carbohydrate, energy, fat, fiber (10%), mineral, protein, and vitamin content. Dry diets were supplemented with fruits, vegetables, and other edible foods (e.g., nuts, cereals, seed mixtures, yogurt, peanut butter, popcorn, marshmallows, etc.) to provide variety to the diet and to inspire species-specific behaviors such as foraging. To further promote psychological well-being, animals were provided with food enrichment, human-to-monkey interaction, structural enrichment, and manipulanda. Environmental enrichment objects were selected to minimize chances of pathogen transmission from one animal to another and from animals to care staff. While on study, all animals were evaluated by trained animal care staff at least twice daily for signs of pain, distress, and illness by observing appetite, stool quality, activity level, and physical condition. Animals exhibiting abnormal presentation for any of these clinical parameters were provided appropriate care by attending veterinarians.

### Care & Use of Macaques

The female macaques described in this report were co-housed with a compatible male and observed daily for menses and breeding. Pregnancy was detected by abdominal ultrasound, and gestational age was estimated as previously described [26]. For physical examinations, virus inoculations, and blood collections, dams were anesthetized with an intramuscular dose of ketamine (10 mg/kg). Blood samples from the femoral or saphenous vein were obtained using a vacutainer system or needle and syringe. Pregnant macaques were monitored daily prior to and after inoculation to assess general well-being and for any clinical signs of infection (e.g., diarrhea, inappetence, inactivity, and atypical behaviors).

### Viral inoculation and monitoring

Thirteen pregnant dams were inoculated subcutaneously with 10^4^ PFU of ZIKV/*Aedes africanus*/SEN/DAK-AR-41524/1984 (ZIKV-DAK, Genbank accession number KY348860). This virus was originally isolated from *Aedes africanus* mosquitoes with a round of amplification on *Aedes pseudocutellaris* cells, followed by amplification on C6/36 cells and two rounds of amplification on Vero cells. ZIKV-DAK was obtained from BEI Resources (Manassas, VA). Virus stocks were prepared by inoculation onto a confluent monolayer of C6/36 mosquito cells. Inocula were prepared from the viral stock described above. The stock was thawed, diluted in sterile PBS to 10^4^ PFU/mL, and loaded into a 1-mL syringe that was kept on ice until inoculation. Animals were anesthetized as described above, and 1 mL of the inoculum was delivered subcutaneously over the cranial dorsum. Animals were monitored closely following inoculation for any signs of an adverse reaction. Eight pregnant dams served as controls and were inoculated subcutaneously with 1 mL of sterile PBS, using the same procedure as described above. All control animals were subjected to the same sedation and blood collection schedule as the ZIKV inoculated animals.

### Pregnancy termination and tissue collection

Nine of the pregnant dams inoculated with 10^4^ PFU of ZIKV-DAK and all eight pregnant dams inoculated with PBS had their pregnancies surgically terminated at gd 36–46 via laparotomy. This procedure was terminal for two control dams 30/7-C1B and 30/7-C2; and three ZIKV-DAK inoculated dams 14-4, 14-3, and 14-2; because of animal protocol regulations that were not related to viral infection or any particular outcomes of this study. Thus, plasma viremia and plaque reduction neutralization test (PRNT) data points are not provided for 14-4, 14-3, and 14-2 beyond 14 dpi. During the laparotomy procedure, the entire conceptus (fetus, placenta, fetal membranes, umbilical cord, and amniotic fluid) was removed. The tissues for all animals were dissected using sterile instruments which were changed between each organ/tissue to minimize possible cross-contamination. Each organ/tissue was evaluated grossly *in situ*, removed with sterile instruments, placed in a sterile culture dish, and dissected for histology, viral burden assay, or banked for future assays. Samples of the MFI included full-thickness center-cut sections of the primary and secondary placental discs containing decidua basalis and chorionic plate. Biopsy samples were collected for RT-qPCR evaluation including: decidua basalis dissected from the maternal surface of the placenta, chorionic plate, chorionic membrane and amnion. If decidua parietalis was adherent to the chorionic membrane it was removed from the sample collected for RT-qPCR. Fetal tissues and fluids including cerebral spinal fluid (CSF), fetal brain, eye, heart or fetal chest, fetal limb containing muscle and skin, liver, kidney, and spleen were collected for viral burden assay or banked for future assays. The remaining fetal tissues were preserved for histological evaluation.

### Full body PFA perfusion and collection of gravid uterus

Four pregnant dams, 03-1, 06-1, 09-1, and 10-1, were euthanized 10, 9, 6, and 3 days post-maternal inoculation (gd 39, 42, 36, and 32), respectively. Euthanasia was performed by first anesthetizing the animal with an intramuscular dose of ketamine (at least 15 mg/kg) followed by an intravenous overdose (greater than or equal to 50 mg/kg or to effect) of sodium pentobarbital. A terminal full-body perfusion (through the heart) with 4% PFA was then performed. Following perfusion, the intact, gravid uterus was removed. The uterus was shallowly scored on the dorsal/posterior side coronally and on ventral/anterior side longitudinally to annotate orientation. The uterus was then placed in 4% PFA with a stir bar on a magnetic stir plate for 24 hours. After the initial 24 hours, the cervix was removed to expose the internal cervical os. Similarly, the serosa and myometrium of the fundus were excised to the level of the endometrium. This increased the 4% PFA penetration for optimal fixation. The uterus was placed in fresh 4% PFA with a stir bar for an additional 48 –72 hours. After a total of 72 – 96 hours of fixation in 4% PFA, the entire uterus was placed in a sectioning jig (a slicing box) coronal sectioning. 10-1 and 06-1 were both sliced into 4 mm thick sections, and 03-1 was sliced into 2 mm thick sections. The fetus was detached from the umbilical cord and removed from the uterine lumen during tissue sectioning. The fetus was longitudinally bisected when large enough, or left intact. Coronal sections of the uterus were trimmed into Supa Mega cassettes (Cellpath, Newtown, Mid Wales, United Kingdom) (Fig. 1). The fetus was placed in the center of an intact middle slice of the gravid uterus. Cassettes were fixed in 4% PFA for an additional 12 hours before changing to 70% ethanol for routine processing and paraffin embedding for histology and unstained sections were cut using a microtome.

### Viral RNA isolation from blood, tissues, and other fluids

RNA was isolated from maternal and fetal plasma, CSF, and amniotic fluid using the Viral Total Nucleic Acid Purification Kit (Promega, Madison, WI) on a Maxwell 48 RSC instrument (Promega, Madison, WI) as previously reported [54]. Fetal and maternal tissues were processed with RNAlater (Invitrogen, Carlsbad, CA) according to the manufacturer’s protocols. RNA was recovered from tissue samples using a modification of a previously described method [55]. Briefly, up to 200 mg of tissue was disrupted in TRIzol (Life Technologies, Carlsbad, CA) with 2 x 5 mm stainless steel beads using a TissueLyser (Qiagen, Germantown, MD) for 3 minutes at 25 r/s for 2 cycles. Following homogenization, samples in TRIzol were separated using Bromo-chloro-propanol (Sigma-Aldrich, St. Louis, MO). The aqueous phase was collected, and glycogen was added as a carrier. The samples were washed in isopropanol and ethanol precipitated. RNA was re-suspended in 5 mM Tris pH 8.0 and stored at -80 °C. RNA isolated using this method was used for the quantification of ZIKV RNA via RT-qPCR and for the detection IFNλ-1 mRNA in the chorionic membrane samples.

### RNA isolation from placenta biopsies for IFNλ-1 evaluation

Placenta biopsies were collected and preserved in RNA later as described above. Biopsies were homogenized and the RNA isolated using methods previously described [56]. Briefly, preserved tissue was disrupted in TRIzol (Invitrogen, Waltham, MA) with 2 x 3.2 mm stainless steel beads using a TissueLyser (Qiagen, Germantown, MD) for 3 minutes at 30 r/s for 3 cycles. Samples were then spun down, and supernatant was collected. Homogenized samples were kept in TRIzol at -80 °C until RNA extraction. The TRIzol-cell mixture was layered onto a Phasemaker Tube (Invitrogen, Waltham, MA) and incubated at room temperature for 3 min followed by the addition of 200 µL of chloroform (MP Biomedicals, Santa Ana, CA) and spun at 16,000 x g at 4 °C for 15 min to allow separation. The aqueous layer was recovered, 500 µL 70% ethanol was added, and then transferred a RNeasy Mini Kit (Qiagen, Germantown, MD) spin column and proper buffer washing steps were carried out following manufacturer recommendations including a 15 min on-column DNAse (Qiagen, Germantown, MD) treatment. Quantification and quality assessment of total RNA was performed using a NanoDrop Spectrophotometer (Thermo Scientific, Waltham, MA).

### Quantitative reverse transcriptase (RT-qPCR) for ZIKV RNA

ZIKV RNA was isolated from both fluid and tissue samples as previously described [39]. Viral RNA was then quantified using a highly sensitive RT-qPCR assay based on the one developed by Lanciotti et al. [57], though the primers were modified with degenerate bases at select sites to accommodate African-lineage Zika viruses. RNA was reverse-transcribed and amplified using the TaqMan Fast Virus 1-Step Master Mix RT-qPCR kit (Life Technologies, Carlsbad, CA, USA) on the LC96 instrument (Roche, Indianapolis, IN, USA), and quantified by interpolation onto a standard curve made up of serial tenfold dilutions of *in vitro* transcribed RNA. RNA for this standard curve was transcribed from a plasmid containing an 800 bp region of the Zika virus genome that is targeted by the RT-qPCR assay. The final reaction mixtures contained 150 ng random primers (Promega, Madison, WI), 600 nM each primer and 100 nM probe. Primer and probe sequences are as follows: forward primer: 5’-CGYTGCCCAACACAAGG-3’, reverse primer: 5′-CCACYAAYGTTCTTTTGCABACAT-3′ and probe: 5′-6-carboxyfluorescein-AGCCTACCTTGAYAAGCARTCAGACACYCAA. The limit of detection for fluids (plasma, extraembryonic coelomic fluid, amniotic fluid, CSF) with this assay is 150 copies/ml. The theoretical limit of detection for the tissues is 3 copies/mg [23].

### RT-qPCR for type III interferon gene IFNλ-1

Total RNA was reverse transcribed using a SuperScript™ III First-Strand Synthesis SuperMix kit (Invitrogen, Waltham, MA)) following the manufacturer’s recommended protocol and a starting input of 3 μg of total RNA. cDNA was diluted with water 1:5 and RT-qPCR reactions were prepared by combining iQ SYBR green supermix (mBio-Rad, Hercules CA), primers and water as previously described [19]. The primer sequences were as follows: IFNλ-1 forward 5’-ATCGTGGTGCCTGGTGACT TT-3’ and reverse 5’-TTGAGTGACTCTTCCAAGGCA-3’ and beta-actin forward 5’-CTACCATGAGCTGCGTGTGG-3’ and reverse 5’-GTACCATGGCTGGGGTGTTGA-3’. All reactions were performed in triplicate and no template control reactions were performed for each primer. The RT-qPCR reactions were performed as recommended by the kit protocol and run on a Roche Light Cycler (Roche, Indianapolis, IN, USA). The limit for cycle of quantification was defined at less than 29. Beta-actin served as the reference gene for all samples to calculate the ΔCq [58].

### Infectious viral quantification by plaque assay

Extraembryonic coelomic fluid and amniotic fluid samples were evaluated for infectious ZIKV by plaque assay. These samples were selected based on RT-qPCR results showing a high level of ZIKV RNA. Titrations for replication competent virus quantification of amniotic fluid were completed by plaque assay on Vero cell cultures as described previously [24]. Vero cells were obtained from the American Type Culture Collection (CCL-81). Duplicate wells were infected with 0.1 mL of aliquots from serial 10-fold dilutions in growth media and virus was adsorbed for 1h. Following incubation, the inoculum was removed, and cell monolayers were overlaid with 3 mL containing a 1:1 mixture of 1.2% oxoid agar and DMEM (Gibco, Carlsbad, CA, USA) with 10% (vol/vol) FBS and 2% (vol/vol) penicillin/streptomycin. Cells were incubated at 37°C in 5% CO_2_ for 32 days for plaque development. Cell monolayers were then stained with 3 mL of overlay containing a 1:1 mixture of 1.2% oxoid agar and DMEM with 2% (vol/vol) FBS, 2% (vol/vol) penicillin/streptomycin and 0.33% neutral red (Gibco). Cells were incubated overnight at 37°C and plaques were counted. Limit of detection for the plaque assay is 0.7 log10 PFU/mL.

### Plaque Reduction Neutralization test (PRNT)

Macaque serum samples were assessed for ZIKV neutralizing antibodies utilizing a plaque reduction neutralization test (PRNT). Endpoint titrations of reactive sera, utilizing a 90% cutoff (PRNT_90_), were performed as previously described [59] against ZIKV/Aedes africanus/SEN/DAK-AR-41524/1984 (ZIKV-DAK). Briefly, ZIKV was mixed with serial 2-fold dilutions of heat-inactivated serum for 1 hour at 37°C before being added to Vero cells, and neutralization curves were generated using GraphPad Prism software (La Jolla, CA). The resulting data were analyzed by nonlinear regression to estimate the dilution of serum required to inhibit both 90% and 50% of infection.

### Histology

Tissues were fixed in 4% PFA, routinely processed, and embedded in paraffin. Paraffin sections were stained with hematoxylin and eosin (H&E) or used for in situ hybridization (ISH), immunohistochemistry (IHC), or immunofluorescence (IF). Histological evaluation of fetal tissues and placentas from the 16 dissected pregnancies was performed by American College of Veterinary Pathologists board-certified pathologists (HAS and PB) blinded to the ZIKV RNA results. Additional histologic examination of placental center cuts from the 16 dissected pregnancies was performed by a placental pathologist (TKM) who was blinded to ZIKV RNA results and pregnancy treatment but was informed of the gestational age at the time of collection to appropriately evaluate villous maturation. Full thickness placental center cuts were collected from the center of each placenta and contained the decidua, placental villi, chorionic plate. Histologic data from some of the control animals (30/7-C1, 30/14-C1, 30/14-C2, and 30/14-C3) were also used for a different study [15] contemporary with this one. Uteroplacental histologic sections were scored for the presence or absence of decidual vasculitis, chronic villitis, remote (chronic) infarction, acute infarction, and villous calcification. Evaluation of the four intact gravid uteri was done by TKM in an unblinded fashion, with the treatment and tissue viral burden known to the pathologist. Brightfield photomicrographs were taken using the Nikon Eclipse Ti2 (Nikon Instruments Inc., Melville, NY, USA) using NIS-Elements AR software version 5.02.006 (Nikon Instruments Inc., Melville, NY, USA) or on Olympus BX46 (Olympus, Center Valley, PA, USA). Additional brightfield photomicrographs were obtained with an Aperio Digital Pathology Slide Scanner (Leica Biosystems, Deer Park, IL, USA). Scale bars were added using NIS-Elements AR, and photomicrographs were white-balanced using Adobe Photoshop 2020 version 21.10 (Adobe Inc, San Jose, CA, USA).

### Detection of ZIKV RNA using *in situ* hybridization (ISH)

ISH was conducted as previously described [26]. The ISH probes against the Zika virus genome were purchased commercially (Advanced Cell Diagnostics, Cat No. 468361, Newark, California, USA). Tissue sections were determined to be negative when ZIKV RNA was undetectable by RNAscope ISH, including instances when only a single, obviously non-cellular, focus of staining was observed. Tissue sections were determined to be positive for ZIKV RNA as detectable by RNAscope ISH when there was clear staining within specific tissues and anatomic structures. Any positive signal that was limited to a single cell-sized focus of indeterminate histological appearance (specks) of positive signal was interpreted as non-specific staining and not interpreted as positive. Uninfected tissue sections served as negative controls (S18).

### Detection of ZIKV replication using multiplex fluorescence *in situ* hybridization (mFISH)

Multiplex fluorescence in situ hybridization (mFISH) was performed using the RNAscope® Fluorescent Multiplex Kit (Advanced Cell Diagnostics, Newark, CA) according to the manufacturer’s instructions with modifications. Briefly, twenty ZZ probe pairs with C1 channel (green, Cat# 463781) targeting ZIKV positive sense RNA and forty ZZ probe pairs with C3 channel (red, Cat# 467911) targeting ZIKV negative sense RNA were synthesized by Advanced Cell Diagnostics. Paraformaldehyde-fixed paraffin-embedded tissue sections were deparaffinized, followed by antigen retrieval. Sections were exposed to ISH target probes and incubated at 40°C in a hybridization oven for two hours. After rinsing, ISH signal was amplified using company-provided Pre-amplifier and Amplifier conjugated to fluorescent dye. Sections were counterstained with 4’, 6-diamidino-2-phenylindole (DAPI, Thermo Fisher Scientific, Waltham, MA, USA), mounted, and stored at 4°C until image analysis. FISH images were captured on an LSM 880 Confocal Microscope with Airyscan (Zeiss, Oberkochen, Germany) and processed using open-source ImageJ software (National Institutes of Health, Bethesda, MD, USA). Uninfected tissue sections served as negative controls.

### Immunohistochemistry

Paraffin sections were deparaffinized and subjected to a heat induced antigen retrieval protocol (10 mM sodium citrate with 0.05% Tween-20 (Fisher BioReagents, Waltham,

MA, USA) at 110°C for 15 min). Slides were then blocked with 3% hydrogen peroxide for 10 minutes and with Background Punisher (Biocare Medical, Concord, CA, USA) for 10 minutes. Sections were immunohistochemically stained using a proprietary polymer-based peroxidase staining method (Biocare Medical, Concord, CA, USA) and incubated with primary antibodies (S3 table) overnight at 4°C. Slides were processed for bound antibody detection by incubation in full-strength MACH2 Polymer-horseradish peroxidase conjugate (Biocare Medical, Concord, CA, USA) for 20 min and developed with Betazoid Diaminobenzidine (DAB) chromogen (Biocare Medical, Concord, CA, USA). All washes were in tris-buffered saline pH 8.4 with Tween-20 (Fisher BioReagents, Waltham, MA, USA) at room temperature.

### Immunofluorescence

Tissue section slides were prepared from paraformaldehyde-fixed, paraffin-embedded blocks. Slides were deparaffinized in xylene and hydrated through an ethanol series. Antigen retrieval was performed in a citrate buffer (pH 6; 10mM sodium citrate with 0.05% Tween 20) via microwave for 5 min at 95°C. Slides were then blocked in a blocking buffer (2% goat serum, 1% bovine serum albumin, 0.1% Triton X100 and 0.05% Tween-20 in tris buffered saline) at room temperature for one hour. Primary antibodies (S3 table) were diluted in a blocking buffer and incubated on the slides overnight at 4° C. The following day, slides were incubated with secondary antibodies (S3 table) for 1 hour at room temperature. Slides were coverslipped with a DAPI inclusive mounting media (Prolong Anti Fade, Thermofisher, Waltham MA, USA). Stained tissue sections were imaged using an EVOS AutoFL microscope system (Life Technologies, Grand Island, NY, USA). The light cubes used for fluorescent imaging were Texas Red, Cy5 and DAPI (Life Technologies, Grand Island, NY, USA) for detecting ALEXA 594, ALEXA 647 and DAPI, respectively. Imaging was processed using open-source ImageJ software (National Institutes of Health, Bethesda, MD, USA). Control staining was done with isotype controls detailed in S3 table, imaging of the control staining can be found in S19 Fig.

### Estradiol and progesterone measurement with Roche cobas e411

Serum samples were analyzed for estradiol and progesterone using a cobas e411 analyzer equipped with ElectroChemiLuminescence technology (Roche, Basal, Switzerland) according to manufacturer instructions. Results were determined via a calibration curve which was instrument-generated by 2-point calibration using traceable standards and a master curve provided via the reagent barcode. Inter-assay coefficient of variation (CV) was determined by a pool of rhesus plasma. For estradiol, the limit of quantitation (LOQ) was 25 pg/mL, the intra-assay CV was 2.02%, and the inter-assay CV was 5.05%. For progesterone, the LOQ was 0.2 ng/mL, the intra-assay CV was 1.37%, and the inter-assay CV was 4.63%. Serum hormone levels were evaluated by first calculating the area under the curve for estradiol and progesterone for each dam using GraphPad Prism software (La Jolla, CA). Area under the curve was compared between ZIKV-infected dams and controls using Welch’s t-test using GraphPad Prism software (La Jolla, CA).

## Acknowledgments

We thank the Wisconsin National Primate Research Center (WNPRC) Veterinary, Scientific Protocol Implementation, Pathology Service Unit, and Behavioral Management staff for assistance with animal procedures, including breeding, monitoring, surgery, and necropsy. We thank the WNPRC Virology service and assay service units for performing viral RNA isolation, ZIKV RNA qRT-PCR, and the hormone assays. We thank the University of Wisconsin Translational Research Initiatives in Pathology (TRIP) Laboratory, supported by the UW Department of Pathology and Laboratory Medicine, UWCCC (P30 CA014520) and the Office of Research Infrastructure Programs-NIH (S10 OD023526) for use of its facilities and services.

## Supporting information captions

**S1 Fig. Plasma viremia of dams inoculated with 10^4^ PFU of ZIKV-DAK.**

Plasma viremia was determined by RT-qPCR and is represented as vRNA copies/mL from 0 up to 16 dpi. Only values above the limit of detection (LOD) of 150 copies/mL are shown. LOD is represented on the graph with a dashed line. (A) Plasma viremia for the eight animals in the fetectomy cohort 1. (B) Plasma viremia for the four animals in the perfusion cohort.

**S2 Fig. Neutralizing antibody titers pre-and post-inoculation with 10^4^ PFU of ZIKV-DAK in the fetectomy cohort dams.**

Plaque reduction neutralization tests (PRNT) were performed on serum samples collected prior to infection and at 14 or approximately 28 dpi. Data are expressed relative to infectivity in the absence of serum. A) Neutralization curves. B) PRNT_90_ and PRNT_50_ values were estimated using nonlinear regression analysis and are indicated with dotted lines in Panel A.

**S3 Fig. Neutralizing antibody titers pre- and post-inoculation with 10^4^ PFU of ZIKV-DAK for the perfusion cohort dams.**

Plaque reduction neutralization tests (PRNT) were performed on serum samples collected prior to infection and on the day of euthanasia. PRNTs are expressed relative to infectivity in the absence of serum. All dams had no neutralizing antibodies prior to inoculation. Symbols for each dam is shown in the right side of the figure, neutralizing antibodies were not detected in any of the dam’s 0 dpi samples.

**S4 Fig. ZIKV RNA detection in additional fluids.**

Viral load was determined by RT-qPCR and is represented as vRNA copies/mL. The dashed lines represent the LOD of 150 copies/mL. (A) ZIKV RNA burden in the extraembryonic coelomic fluid at 7 dpi. (B) ZIKV RNA burden in the umbilical cord blood at 7 dpi. (C) ZIKV RNA burden in the extraembryonic coelomic fluid at 14 dpi. (D) ZIKV RNA burden in the amniotic fluid at 14 dpi.

**S5 Fig. Replicating ZIKV detected in the chorionic membrane at 14 dpi.** Photomicrographs of multiplex fluorescence in situ hybridization (mFISH) to detect genomic, positive sense ZIKV RNA (green), and replicative intermediate negative sense RNA (red) with nuclear DAPI staining (blue), colocalization of positive and negative sense ZIKV RNA is seen as yellow. (A)(C)(E)(G) panels show merged images of all channels (red, green, and blue). (B)(D)(F)(H) shows only the red channel showing the replicative intermediate RNA detected of the panel immediately to the left. (A)(B)(C)(D) are from case 14-3. (E)(F)(G)(H) are from case 14-4. (A)(C) 50 µm (E) and (G) 20 µm.

**S6 Fig. Chorionic plate infection at 14 dpi.**

(A) Full thickness decidual/placental tissue section with ISH detection of ZIKV RNA shown as pink staining. The scale bar represents 1000 µm. The Orange square indicates the area magnified in (B). (C) corresponding section H&E stained. (D) Photomicrograph of IF staining for CD163 (red) ZIKV (green) and DAPI (blue) in the chronic plate in a subsequent section of the slide shown in (A) Dashed circle identifies cells that are positive for both CD163 and ZIKV. The scale bar represents 200 µm. (B)(C) Scale bars represent 100 µm.

**S7 Fig. ZIKV infection of the amnion at 14 dpi.**

(A) Pink staining indicates ZIKV RNA detected via ISH. Orange square indicates the area magnified in (C). (B) H&E stained section. Red square indicates the area magnified in (D). (A)(B) The scale bars represent 100 µm. (C)(D) The scale bars represent 50 um. (E) IF staining for CD163 (red) ZIKV (green) and DAPI (blue). Little to no colocalization of ZIKV and CD163 was seen as indicated by arrows. (F) IF staining for cytokeratin (red) to identify amniotic epithelial cells, ZIKV (green), and DAPI (blue). Little cytokeratin staining was seen. Positive cytokeratin staining is indicated by arrows and no colocalization of ZIKV and cytokeratin. (E)(F) Scale bars represent 200 µm. (G) Multiplex fluorescence in situ hybridization (mFISH) to detect genomic, positive sense ZIKV RNA (green), and replicative intermediate negative sense RNA (red) with nuclear DAPI staining (blue), colocalization of positive and negative sense ZIKV RNA is seen as yellow. The scale bar represents 50 µm. (H) The red channel of the photomicrograph shown in (G) shows only the replicative intermediate RNA.

**S8 Fig. Replicating ZIKV in the fetus at 14 dpi.**

(A)(B) Multiplex fluorescence in situ hybridization (mFISH) to detect genomic, positive sense ZIKV RNA (green), and replicative intermediate negative sense RNA (red) with nuclear DAPI staining (blue), colocalization of positive and negative sense ZIKV RNA is seen as yellow. Image on the left shows all (red, green, and blue) channels and the image on the left shows the red channel (replicative intermediate RNA) alone. (A) The scale bar represents 20 µm. (B) The scale bar represents 50 µm.

**S9. ZIKV does not reach the gravid uterus at 3 dpi.**

Photomicrograph of a representative coronal section of the gravid uterus evaluated at 3 days post-infection evaluated with ISH. No virus was detected in any of the coronal sections.

**S10 Fig. Additional sites of ZIKV infection in the decidua at 10 dpi.** Photomicrographs of ZIKV RNA shown as pink staining detected via ISH in a representative slide from 09-1. Center image shows the full slide. Color boxes indicate areas of the decidua presented at higher magnification.

**S11 Fig. ZIKV infection of decidua at 10 dpi.**

(A) Coronal section of the uterus evaluated for ZIKV RNA via ISH. The black square indicates the area shown at higher magnification in (B) showing an infected decidual vessel. (C) shows the corresponding H&E staining. (B)(C) The scale bar represents 100 µm.

**S12 Fig. ZIKV infection in the trophoblastic shell in a slice taken from the superior pole of the uterus.**

(A) Full slide with pink staining indicating ZIKV RNA detected via ISH. The scale bar represents 1955 µm. The orange square indicates the portion of the slide shown at higher magnification in (B). Corresponding sections of (B) are IHC stained for cytokeratin shown as brown staining in (C) and H&E stained in (D).

**S13 Fig. ZIKV infection of endothelial cells in chorionic plate vessels.**

(A) Full slide of ISH detection of ZIKV RNA shown as pink staining. The orange square indicates the portion of the slide shown at higher magnification in (B). The scale bar represents 50 µm. The yellow square indicates the portion of the slide magnified in (C). The scale bar represents 71 µm. Arrows indicated infected endothelial cells.

**S14 Fig. Replicating virus detected in the placenta and fetus at 10 dpi.**

(A) Full slide of ISH detection of ZIKV RNA shown as pink staining. Colored squares indicate areas of the slide shown at higher magnification in subsequent panels. Yellow (B) shows infection near the chorionic plate, red (C) shows infection in the placental villi, blue (D) and (E) show infection in the fetus. (B)(C)(D)(E) Photomicrographs of multiplex fluorescence in situ hybridization (mFISH) to detect genomic, positive sense ZIKV RNA (green), and replicative intermediate negative sense RNA (red) with nuclear DAPI staining (blue), colocalization of positive and negative sense ZIKV RNA is seen as yellow. The presence of yellow and red staining indicates that there is replicating ZIKV in the chorionic plate, villi, and fetus in this 10 dpi case. The scale bars represent 50 µm.

**S15 Fig. ZIKV infects the peripheral margin of the placenta and trophoblasts in the chorionic plate.**

(A) Full slide evaluated with ISH for ZIKV RNA indicated by pink staining. Black square indicated the portion of the section magnified in (B) (C) Corresponding H&E stained section. PM = the peripheral margin of the placenta, CM = the chorionic membrane and CP = chorionic plate. The pink staining in the chorionic plate includes trophoblasts of the chorionic plate.

**S16 Fig. Serum levels of estradiol and progesterone.**

(A)(C)(E)(G) Mean serum levels of estradiol (A)(E) and progesterone (C)(G). Red dots represent mean values for ZIKV-infected dams, black dots represent mean values for mock-infected dams. Ticks represent the range of values for each group. (A)(C) represent the values 0–7 dpi for all eight ZIKV-infected dams from cohort 1 and all eight mock-infected dams. (E)(G) show the values for only the four dams from the 14 dpi group in cohort 1 and their respective controls (n=4) for serum hormone levels 0–14 dpi. (B)(D)(F)(H) Calculated area under the curve for respective serum hormone levels. Area under the curve of the ZIKV-infected was compared to the mock-infected using Welch’s t-test. There were no significant differences between groups (ns). (B)(D) ZIKV-infected dams (n=8) are compared to mock-infected dams (n=7). One dam had two pregnancies randomly assigned to the 7 dpi mock group. Therefore, an average value between her two pregnancies was used in (B) and (C); thus only seven data points are used for the mock-infected dams. (A)(E) Limit of quantification for the serum estradiol assay was 25 pg/mL and is represented by the dashed line. (C)(G) Limit of quantification for the serum progesterone assay was 0.2 ng/mL and is represented by the dashed line.

**S17 Fig. Dam 14-5 did not develop plasma viremia or expand neutralizing antibody titers post-inoculation with 10^4^ PFU of ZIKV-DAK.**

(A) Plasma ZIKV loads were determined by RT-qPCR and are represented as vRNA copies/mL from 0 up to 16 dpi. LOD is represented on the graph with a dashed line. (B)(C) Plaque reduction neutralization tests (PRNT) were performed on serum samples collected prior to infection and 27 dpi. Data are expressed relative to infectivity in the absence of serum. (B) Neutralization curves. (C) PRNT_90_ and PRNT_50_ values were estimated using nonlinear regression analysis and are indicated with dotted lines in Panel A.

**S18 Fig. Uninfected Control Tissue ZIKV RNA detection via ISH.** Photomicrographs showing different regions of a placenta from an uninfected control pregnancy. (A) Placental villi with decidua, (B) Chorionic plate with placental villi.

**S19. IF isotype control staining images.**

**S1 Table. Details on each rhesus macaque pregnancy. S2 Table. Histopathology results.**

**S3 Table. Antibody information.**

